# Integrative functional genomics decodes herpes simplex virus 1

**DOI:** 10.1101/603654

**Authors:** Adam W. Whisnant, Christopher S. Jürges, Thomas Hennig, Emanuel Wyler, Bhupesh Prusty, Andrzej J Rutkowski, Anne L’hernault, Margarete Göbel, Kristina Döring, Jennifer Menegatti, Robin Antrobus, Nicholas J. Matheson, Florian W.H. Künzig, Guido Mastrobuoni, Chris Bielow, Stefan Kempa, Liang Chunguang, Thomas Dandekar, Ralf Zimmer, Markus Landthaler, Friedrich Grässer, Paul J. Lehner, Caroline C. Friedel, Florian Erhard, Lars Dölken

## Abstract

Since the genome of herpes simplex virus 1 (HSV-1) was first sequenced more than 30 years ago, its predicted 80 genes have been intensively studied. Here, we unravel the complete viral transcriptome and translatome during lytic infection with base-pair resolution by computational integration of multi-omics data. We identified a total of 201 viral transcripts and 284 open reading frames (ORFs) including all known and 46 novel large ORFs. Multiple transcript isoforms expressed from individual gene loci explain translation of the vast majority of novel viral ORFs as well as N-terminal extensions (NTEs) and truncations thereof. We show that key viral regulators and structural proteins possess NTEs, which initiate from non-canonical start codons and govern subcellular protein localization and packaging. We validated a novel non-canonical large spliced ORF in the ICP0 locus and identified a 93 aa ORF overlapping ICP34.5 that is thus also deleted in the FDA-approved oncolytic virus Imlygic. Finally, we extend the current nomenclature to include all novel viral gene products. Taken together, this work provides a valuable resource for future functional studies, vaccine design and oncolytic therapies.

## Main

Herpes simplex virus 1 (HSV-1) is the causative agent of the common cold sores but also responsible for severe, life-threatening disease including generalized skin infections, pneumonia, hepatitis and encephalitis^1^. The HSV-1 genome is about 152kb in size and known to encode at least 80 open reading frames (ORFs), many of which have been extensively studied. Large-scale RNA-seq and ribosome profiling recently revealed that the coding capacity of three other herpesviruses, namely human cytomegalovirus (HCMV), Kaposi’s sarcoma-associated herpesvirus (KSHV) and Epstein-Barr Virus (EBV) is significantly larger than previously thought^2–5^. For HCMV and KSHV, in particular, hundreds of new viral gene products were identified. These result from extensively regulated usage of alternative transcription and translation start sites throughout lytic infection. Moreover, these viruses were found to encode hundreds of short ORFs (sORFs) of unknown function. Similar to their cellular counterparts, these may either regulate translation of viral gene products or encode for functional viral polypeptides^6–8^. To date, the majority of novel viral gene products have not been experimentally validated. Furthermore, the lack of a complete annotation and a revised nomenclature severely hampers functional studies.

### Overview on the applied functional genomics approaches

Here, we employed a broad spectrum of unbiased functional genomics approaches and reanalyzed recently published data to comprehensively characterize HSV-1 gene products (Fig. 1). Our analyses of the viral transcriptome includes time-course experiments of *(i)* total RNA-seq and 4sU-seq data^9^, *(ii)* transcription start site (TiSS) profiling using two complementary approaches (cRNA-seq^2^ and dRNA-seq^10^), *(iii)* reanalysis of recently published third-generation sequencing data from the PacBio^11^ and MinlON^12^ platforms, and *(iv)* RNA localization by RNA-seq of subcellular fractions of both wild-type HSV-1 and the deletion mutant of the key viral RNA export factor ICP27^13^. Analyses of the viral translatome includes *(i)* standard ribosome profiling as well as translation start site (TaSS) profiling using *(ii)* Harringtonine and *(iii)* Lactimidomycin. Novel viral ORFs were validated using whole-cell quantitative proteomics and reverse genetics. To make the annotation and all the obtained data readily accessible to the research community, we provide an HSV-1 genome browser software (available at http://software.erhard-lab.de and as Supplementary Software). Thereby, viral gene expression and all data can be visually examined from whole genome to single-nucleotide resolution.

**Figure 1.**
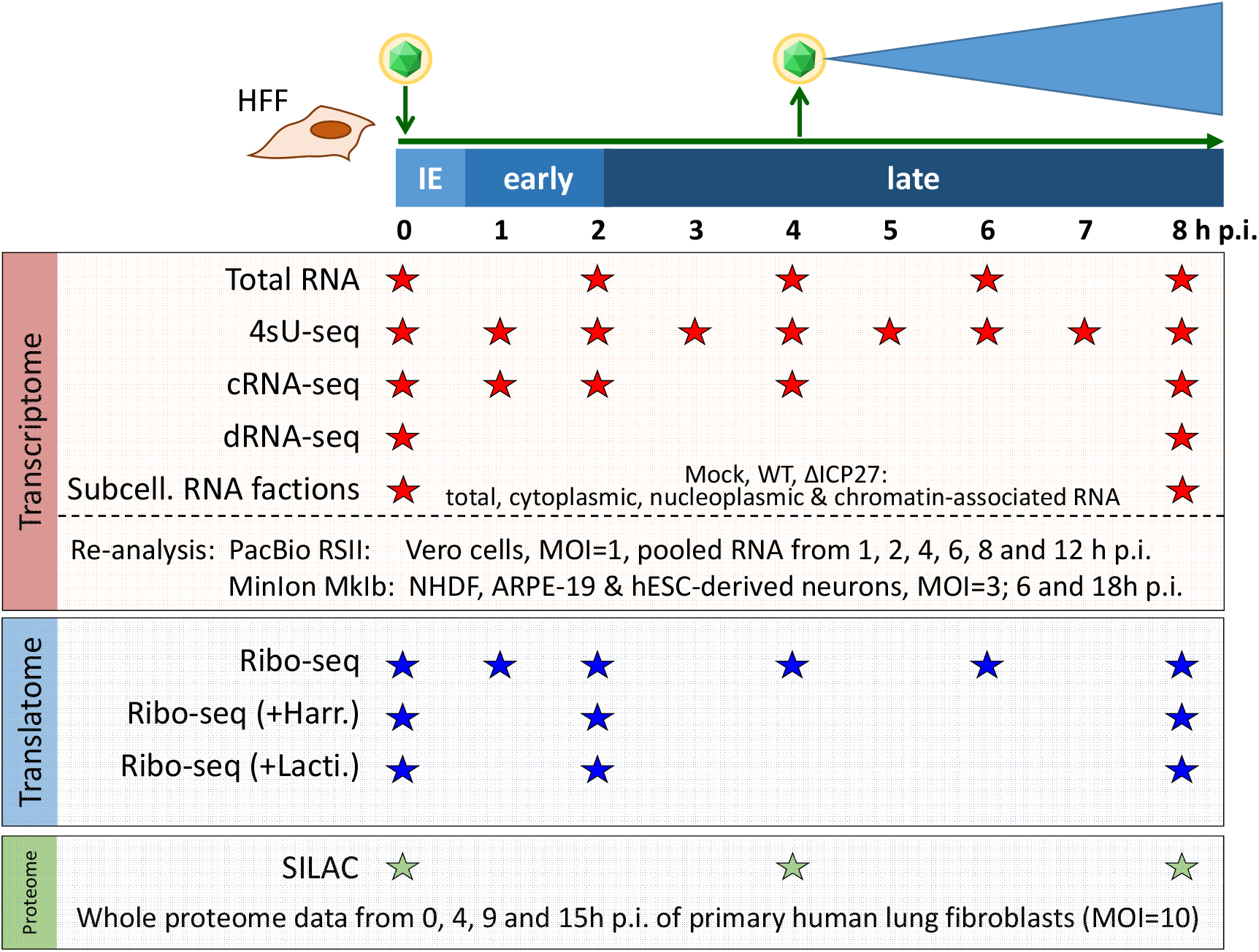
Overview of the applied Omics approaches. Viral gene expression was analyzed in primary human fibroblasts (HFF). The total RNA-seq, 4sU-seq and ribosome profiling data were recently published^9^. To comprehensively identify transcription start site (TiSS), we performed cRNA-seq^2^ and dRNA-seq^10^ as well as RNA-seq on subcellular RNA fractions from mock, wild-type and ΔlCP27 infected cells. Furthermore, we reanalyzed recently published PacBio^11^ and MinlON^12^ sequencing data. Translation start site (TaSS) profiling was performed by ribosome profiling following treatment of cells for 30 min with either Harringtonine or Lactimidomycin^2^. Proteome analysis included two whole proteome data sets using SlLAC and label-free mass spectrometry. The available time points and conditions are indicated by stars.

### Characterization of the HSV-1 transcriptome

To identify the full complement of viral transcripts, we performed TiSS profiling employing a modified RNA sequencing protocol that is based on circularization of RNA fragments (here termed “cRNA-seq”)^2^. It enables quantification of RNA levels as well as identification of 5’ transcript ends by generating a strong enrichment (≈18-fold) of fragments that start at the 5’ RNA ends. This identified 155 TiSS that explained the expression of many previously annotated viral coding sequences (CDS). To comprehensively identify and validate putative novel TiSS, we applied a second 5’-end sequencing approach termed “differential RNA-seq” (dRNA-seq)^10^, which provides a much stronger (≈300-fold) enrichment of TiSS at increased sensitivity. It is based on selective cloning and sequencing of the 5’-ends of cap-protected RNA molecules that are resistant to the 5’-3’-exonuclease Xrn1. The two approaches provided highly consistent data at single nucleotide resolution (Fig. 2a). Furthermore, we reanalyzed recently published third-generation sequencing data obtained using the MinlON^14^ and PacBio^11^ platforms, which confirmed many of the observed TiSS (Suppl. Fig. 1a,b). The 98 viral transcripts recently identified by MinlON generally lacked 7-18 nucleotides (nt) at the 5’ end for technical limitations of the MinlON direct RNA sequencing method (Suppl. Fig. 1b)^15^. Nevertheless, the adjusted TiSS were highly consistent with cRNA-seq and dRNA-seq data (Fig. 2b). Around half of all TiSS previously identified using PacBio sequencing^11^ matched to our data with single nucleotide resolution, the remaining TiSS (104 of 204; 51%) could neither be confirmed by cRNA-seq, dRNA-seq or MinlON (Fig. 2c). Most of them were only called from very few reads and presumably represent cleavage products of larger viral RNAs. ln total, 113 TiSS were identified by at least two of the four approaches. This demonstrates that complementary experiments are essential to exclude false positives and that none of the approaches by themselves is sufficient to reliably identify all viral TiSS.

**Figure 2.**
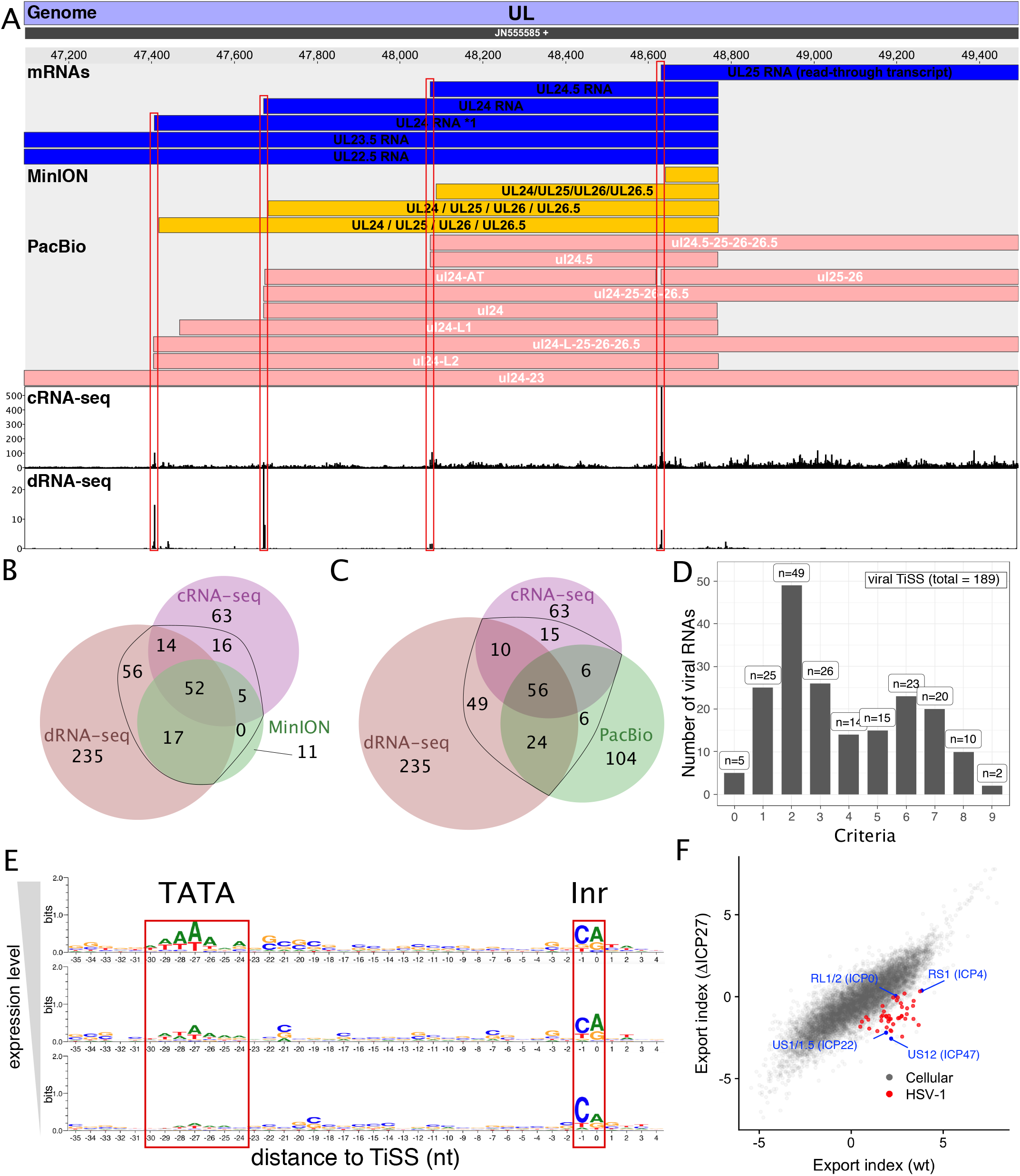
Analysis of viral transcription start sites (TiSS) **(A)** Screenshot of our HSV-1 viewer displaying the annotated mRNAs of MinlON, PacBio and coverage of read 5’-ends for cRNA-seq and dRNA-seq of the UL22.5-UL25 gene locus. Transcripts in our new reference annotation are indicated in blue. **(B,C)** Venn diagram depicting the number of TiSS that were identified by cRNA-seq, dRNA-seq and MinlON **(B)** or PacBio **(C)** sequencing. TiSS included into the final annotation are indicated by the black circle. **(D)** Histogram depicting the number of TiSS criteria that were fulfilled by the individual viral transcripts scored by iTiSS. **(E)** Sequence logos upstream of viral TiSS with viral TiSS grouped into 3 equally sized bins. **(F)** Export index of cellular (grey) and viral (red and blue) gene clusters compared between wild-type HSV-1 (wt) and a null mutant of the viral RNA export factor lCP27 (ΔlCP27). Viral immediate early genes are indicated in blue.

To comprehensively assess the viral TiSS candidates that were only identified by a single approach, we developed a computational approach termed iTiSS (<u>i</u>ntegrative <u>T</u>ranscriptional <u>S</u>tart <u>S</u>ite caller). lt screens putative TiSS candidates deduced from the cRNA-seq, dRNA-seq data, PacBio and MinlON data and scores for differences and temporal changes in read coverage upstream and downstream of a TiSS during the course of HSV-1 infection. ln addition, iTiSS considered transcripts, which were required to explain translation of novel viral ORFs. A single TiSS could thus score up to a total of 9 points. All identified TiSS were manually assessed and curated. ln total, this resulted in 189 viral TiSS of which 159 (84%) were called by at least 2 criteria (Fig. 2d). Three of the five transcripts (LAT^16^, AL-RNA^17^ and US5.1^18^), which we could not confirm by any method, had previously been convincingly validated by other groups and were thus included. The other two were included after careful manual inspection. The complete set of HSV-1 transcripts with their respective scores is provided in Suppl. Tab. 1.

TATA-boxes are a key element of eukaryotic promoters located 25 to 30 bp upstream of the TiSS^19^. They are also prevalent for herpesvirus genes^20^. The presence of a TATA-box or TATA-box-like motif upstream of the viral TiSS strongly correlated with the expression levels of the respective transcripts. For weakly transcribed viral RNAs, the respective motifs were rarely observed (p<10^−6^, Fisher’s exact test). In mammalian cells, the TiSS is marked by the initiator element (Inr), core of which is a pyrimidine-purine (PyPu) dinucleotide^21^. Interestingly, PyPu was also prevalent for the viral TiSS independent of expression levels (Fig. 2e). This provides strong evidence for the TiSS of even the most weakly expressed viral transcripts.

We next looked at splicing within the HSV-1 transcriptome based on our total RNA-seq and 4sU-seq data^9^. This confirmed all 8 well-described splicing events and identified 2 additional tandem acceptor sites (“NAGNAG”)^22^ for the third exon of the ICP0 gene (RL2), as well as for the UL36.6 gene. Recently, Tombácz et al proposed 11 novel splicing events based on PacBio sequencing data^11^. Our data confirmed all of these splicing events. However, only 4 of them occurred at relevant levels (Suppl. Fig. 3 and Suppl. Tab. 2). Two of these explained translation of novel small ORFs (UL40.5 iORF and UL40.7 dORF). Finally, we identified 58 novel putative splicing event sites based on our RNA-seq data (Suppl. Fig. 2 and Suppl. Tab. 2). However, all of these showed substantially lower read coverage than the surrounding exons, indicating that they only represented rare events at best. Therefore, we decided not to include all low abundance splicing events from our new reference annotation. In total, we identified 189 viral TiSS giving rise to at least 201 transcripts and transcript isoforms.

### RNA 3’-end processing and export of viral transcripts

Previous studies reported regulated usage of the 47 viral poly(A) sites during productive infection, which appeared to be mediated or at least influenced by the viral ICP27 protein^23–28^. We recently reported that lytic HSV-1 infection results in a widespread but nevertheless selective disruption of transcription termination of host genes^9^. In contrast to the extensive read-through transcription at host poly(A) sites that we observed by 4-8 h p.i., viral gene expression remained mostly unaffected. Recently published third-generation sequencing data proposed numerous very large viral transcripts spanning multiple viral genes^29^. To address the nature of these transcripts and their role in translation, we performed RNA-seq on subcellular RNA fractions (total RNA, cytoplasmic RNA, nucleoplasmic RNA and chromatin-associated RNA) using both wild-type HSV-1^30^ and a null mutant of the viral RNA export factor ICP27 (ΔICP27). Consistent with the well-characterized role of ICP27 in viral mRNA export^13^, all viral transcripts were more efficiently (≈11-fold) exported to the cytoplasm in wild-type than in ΔICP27 HSV-1 infection (Fig. 2f). Interestingly, this even included the spliced immediate early (lE) genes lCP0 (≈5-fold), lCP22 (≈17-fold) and lCP47 (≈27-fold) as well as the unspliced (lE) lCP4 gene (~11-fold). ln chromatin-associated, nuclear and total cellular RNA, considerable numbers of reads were observed within the first 500 nt downstream of viral poly(A) sites (PAS). However, in the cytoplasmic RNA fraction of infected cells, read levels dropped substantially immediately downstream of the PAS (Fig. 3a). This indicates that reads mapping downstream of PAS reflect mRNA precursors, which remain nuclear and, thus, do not contribute to the translated viral transcriptome. However, for some viral genes, e.g. UL30, UL38 and UL43, considerable numbers of reads that mapped downstream of the respective PAS were present in the cytoplasmic RNA fraction. For the UL30 PAS, this became substantially more prominent late in infection (8 h p.i., Fig. 3b). Furthermore, transcription of UL25, which initiates 107 nt upstream of the UL24 PAS, efficiently bypassed the UL24 PAS already from 2 h p.i. on (Suppl. Fig. 3). The same was observed for UL24.5 which represents an N-terminal truncated isoform of UL24. These data confirm previous findings on differential polyadenylation of selective viral genes during productive infection^23–27^.

**Figure 3.**
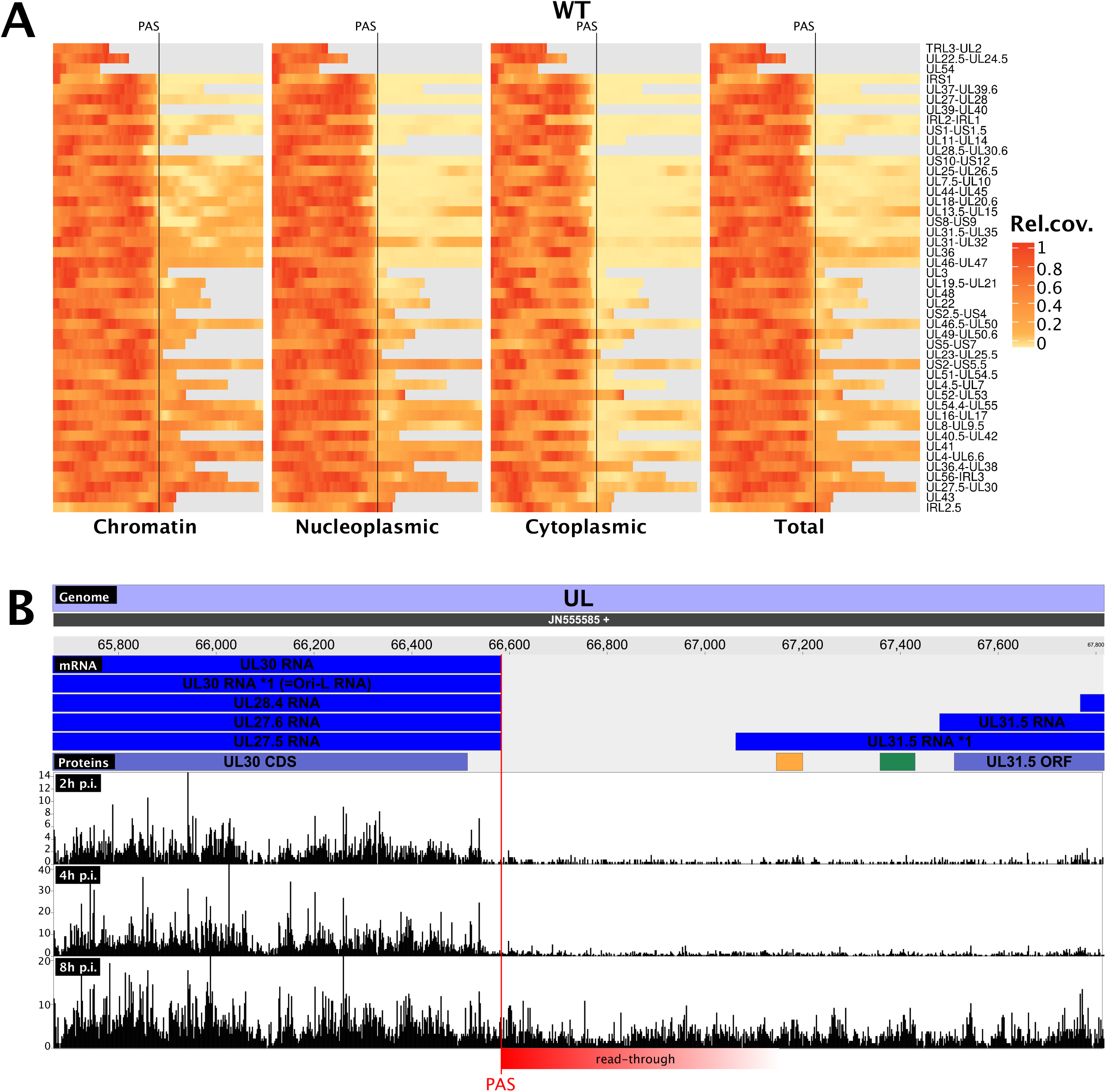
Subcellular localization of viral transcripts. **(A)** Read levels 500 bp upstream (left of PAS indicator) and downstream of the PAS (right of PAS indicator) of viral genes for wild-type HSV-1. Grey bars indicate overlapping parts with other genes, for which reads could not be uniquely assigned. ln the cytoplasmic RNA fraction, read levels dropped substantially immediately downstream of the PAS. **(B)** Screenshot of our HSV-1 viewer depicting poly(A) read-through at the UL30 polyadenylation site (PAS) at 2, 4 and 8 h p.i. of replicate 1. Read-through transcription is indicated in red.

### HSV-1 expresses hundreds of novel ORFs and sORFs

To comprehensively identify the viral translatome, we performed time-course analysis of ribosome profiling as well as translation start site (TaSS) profiling (see Fig. 1 and Supplementary Methods). The obtained data confirmed the expression of all previously annotated ORFs (CDS) but also identified 7 N-terminal truncations (NTTs) and 17 N-terminal extension (NTEs) thereof. Furthermore, we detected 46 novel large ORFs and 134 novel small ORFs (3-99 aa). ln total, we identified 284 viral ORFs (Suppl. Tab. 3a,b). Translation predominantly initiated from AUG start codons (79%). However, non-canonical initiation events also substantially contributed to the HSV-1 translatome with CUG, GUG, ACG and AUC together initiating translation of about 15% and 20% of all large and small viral ORFs, respectively (Fig. 4a,b).

**Figure 4.**
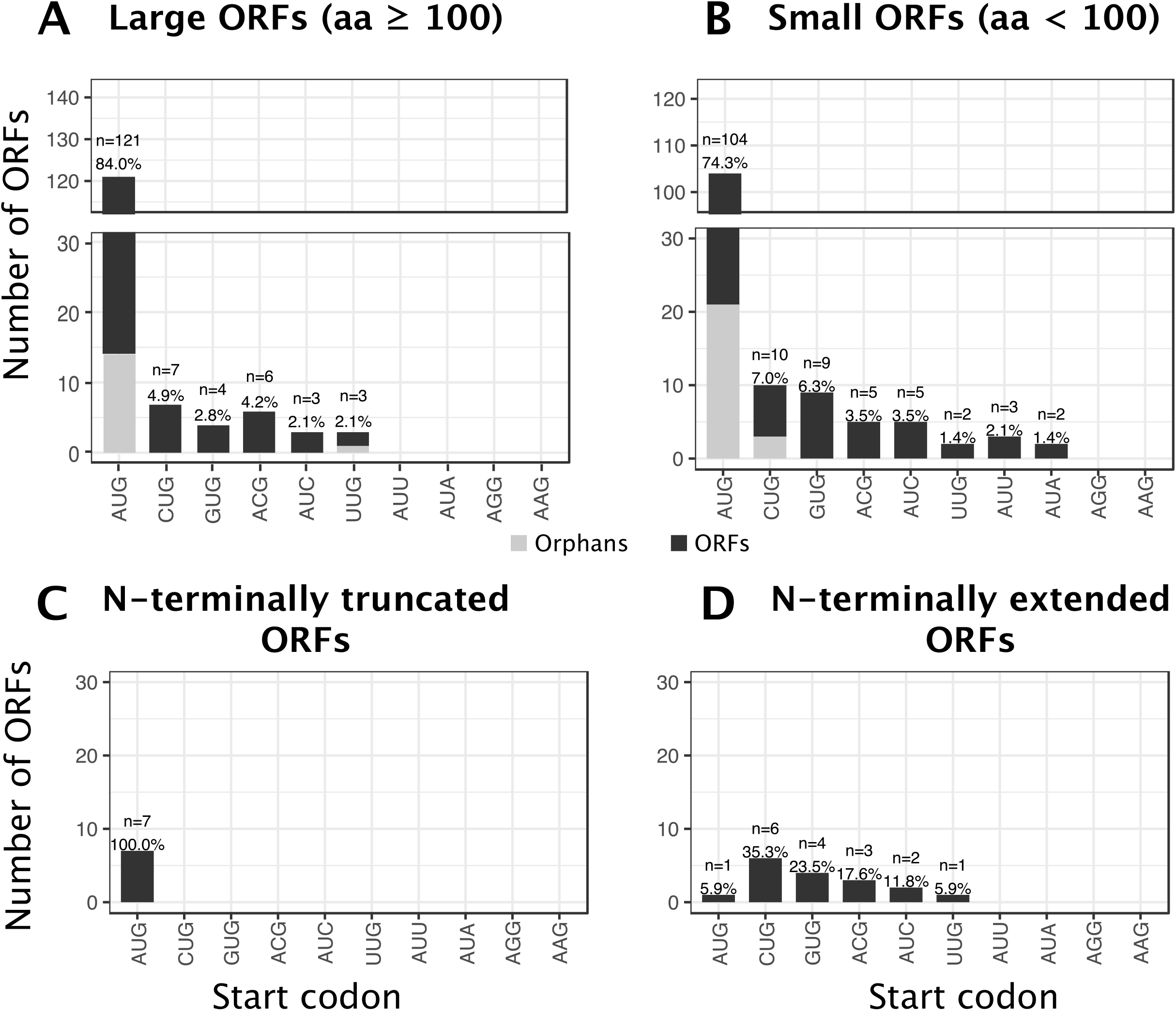
Distribution of start codon usage of all identified HSV-1 proteins. Distribution and frequency of possible start codons used by HSV-1 ORFs **(A)**, sORFs **(B)**, NTTs **(C)** and NTEs **(D)**. Orphan ORFs are depicted in light grey. Six of the previously identified CDS (UL11, UL49.5, US5, US9, US12 and RL2 iso1) are <100 aa and were thus included in **(B)**.

We observed seven NTTs originating from downstream translation initiation events of previously described viral coding sequences (Suppl. Tab. 4). All of these initiated with AUG start codons (Fig. 4c). Alternative TiSS downstream of the main TiSS explained translation of 6 of 7 NTTs (Suppl. Fig. 4). Only for US3.5, we could not identify a corresponding transcript. lt thus remains unclear whether US3.5 is translated from an independent transcript or due to leaky scanning. Six of these NTTs (UL8.5, UL12.5, UL24.5, UL26.5, US1.5 and US3.5) had already been reported^31–36^. Only the NTT of the major DNA-binding protein pUL29 (lCP8; comprising aa 516 to 1212) had so far not been described. lnterestingly, this NTT initiates immediately downstream of the metal-(Zn)-binding loop (residues 499-512)^37,38^. Expression of both a 135 kDa (full-length) and 90kDa (NTT) protein has been shown for the commercial ICP8 antibody 11E2 (Santa Cruz, see product website).

Interestingly, 16 of the 80 viral reference ORFs (20%) showed in-frame NTEs (Suppl. Tab. 5) with translational activity exceeding 10% of the main downstream ORF. The majority of NTEs (16 of 17, including 2 NTEs in UL50) initiated translation from non-AUG start codons (Fig. 4d). This included key viral proteins like the major immediate early protein ICP27, the major capsid protein (VP5, UL19) and the well-studied viral kinase US3. For 5 viral genes, we generated mutant viruses by introducing a triple-FLAG-tag either into the NTE or downstream of the canonical AUG start codon. This confirmed the expression of 6 NTEs including the 2 UL50 NTEs (Fig. 5a-e). Interestingly, the introduction of a triple-FLAG-tag into the N-terminal extension of both ICP27 and VP5 resulted in dead viruses, which could only be reconstituted upon co-transfection of the mutant bacterial artificial chromosomes (BACs) with the wild-type virus BAC. For ICP27, expression of the NTE was already observable when the FLAG-tag was introduced downstream of the canonical AUG-start codon (Fig. 5e). For VP5, the FLAG-tagged NTE appeared to even be dominant negative. Virus reconstitution upon co-transfection with the wild-type HSV-1 BAC resulted in a partial deletion of the NTE within two passages. This indicates that the FLAG-tagged NTE-MCP is assembled into virus particles but renders them dysfunctional due to the N-terminally inserted triple-FLAG-tag.

**Figure 5.**
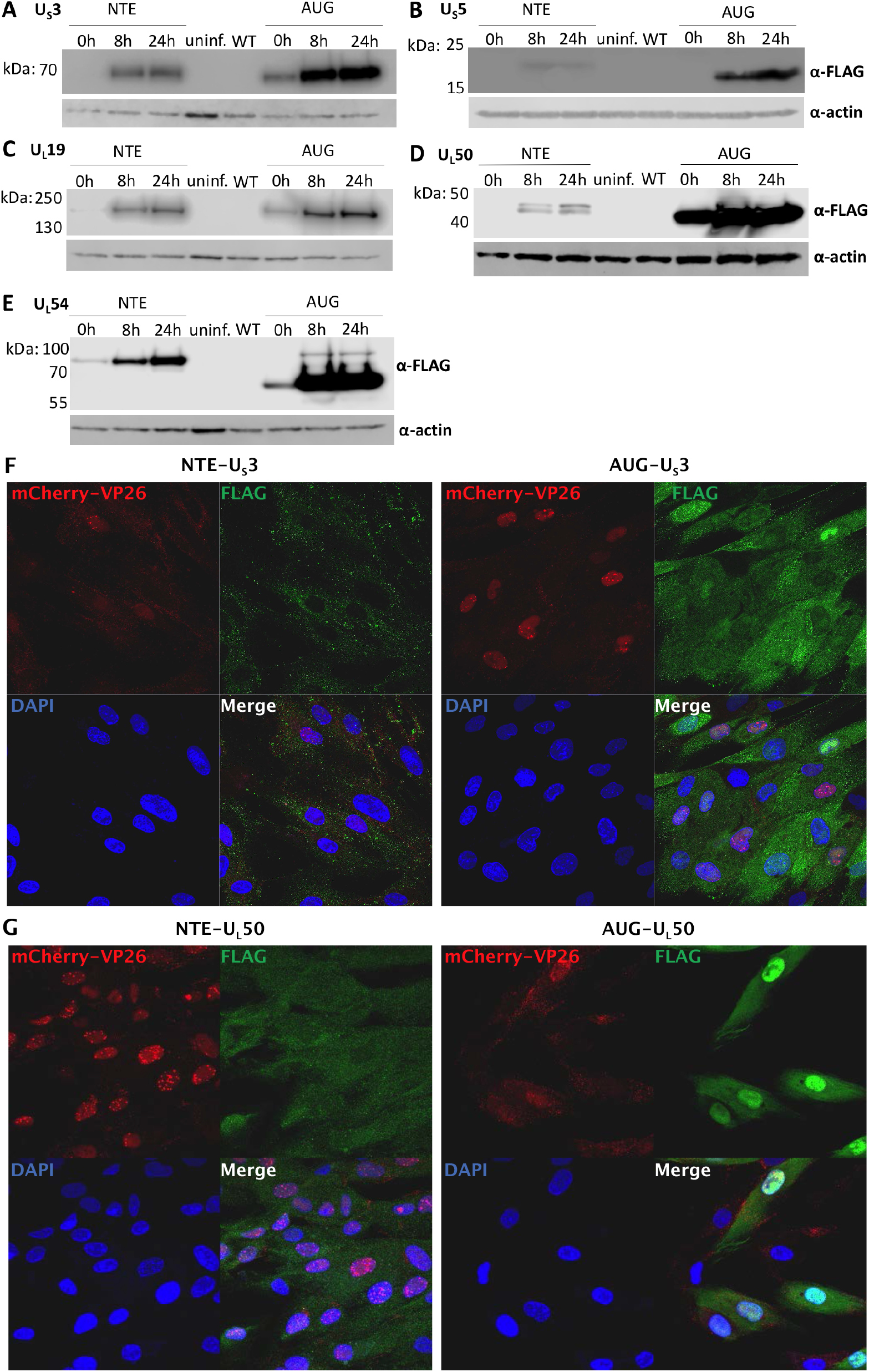
Validation of N-terminal extensions of known HSV-1 proteins. Tagged viruses were generated by inserting a triple FLAG-tag either upstream of the canonical start codon into the N-terminal extension (NTE) or downstream of it (AUG). Western blots of FLAG-tagged N-terminal extensions following infection of human foreskin fibroblasts with the indicated viruses are shown. Expression at the given hours (h) postinfection are compared to uninfected (uninf.) and the parental (WT) virus, both at 24 h p.i. for the HSV-1 genes **(A)** US3, **(B)** US5, **(C)** UL19, **(D)** UL50, and **(E)** UL54. Expression of the NTE of UL54 (ICP27) was already visible when the FLAG-tag was inserted downstream of the canonical AUG. **(F)** Immunofluorescence of human foreskin fibroblasts infected with Vp26-mCherry HSV-1 containing FLAG-tags inserted upstream of the canonical start codon into the N-terminal extension (NTE) or downstream of it (AUG) for US3 and **(G)** UL50. Cell nuclei were stained using DAPI. Protein localization of both NTEs shifts to the cytoplasm. For US3, this is explained by a conserved nuclear localization signal within the NTE.

To test the impact of the respective NTEs on protein localization, we performed immunofluorescence microscopy of both the NTE- and AUG-tagged viruses. While subcellular localization of the NTEs of ICP27 (UL54) and US5 were indistinguishable from their canonical counterparts (Suppl. Fig. 5), the NTEs of US3 and UL50 notably altered subcellular localization (Fig. 5f,g). While canonical US3 was predominantly nuclear, the NTE-US3 localized to the cytoplasm. The US3 NTE contains a leucine-rich stretch indicating a functional nuclear export signal. Pseudorabies virus (PRV), a porcine alphaherpesvirus, expresses two isoforms of US3, both of which initiate from AUG start codons on separate transcripts (Suppl. Fig. 6). The longer isoform encodes a mitochondrial localization signal resulting in the cytoplasmic localization and a failure of the respective protein to be incorporated into the tegument^39^. The DNA sequence of the US3 NTE is conserved in HSV-2 and its role as a nuclear export signal fits data demonstrating that HSV-2 US3 fails to accumulate in the cytoplasm when nuclear export is inhibited^40^. Similar to US3, localization of NTE-UL50 also shifted to the cytoplasm (Fig. 5g). UL50 dUTPase activity in PRV-infected cells was reported to be nuclear^41^, while it appears to be predominantly cytoplasmic with HSV-2^42^ and nearly equally distributed in HSV-1. We conclude that NTEs initiating from non-AUG start codons are common in alphaherpesvirus proteomes. They allow the expression of alternative protein isoforms with different subcellular localization and regulatory motifs.

In 2015, the first oncolytic virus (talimogene laherparepvec (lmlygic)) was approved for therapy of melanoma^43^. This modified herpes simplex virus 1 lacks two viral genes (lCP34.5 and ICP47) and expresses GM-CSF to recruit and stimulate antigen-presenting cells. Within the lCP34.5 (RL1) locus, we found a novel 93 aa ORF, which we termed RL1A (Fig. 6a). lt initiates from an AUG start codon 46 nt upstream of the AUG start codon of RL1 and is translated from the same transcript at >4-fold higher levels. lmlygic thus lacks a third viral protein, namely RL1A.

**Figure 6.**
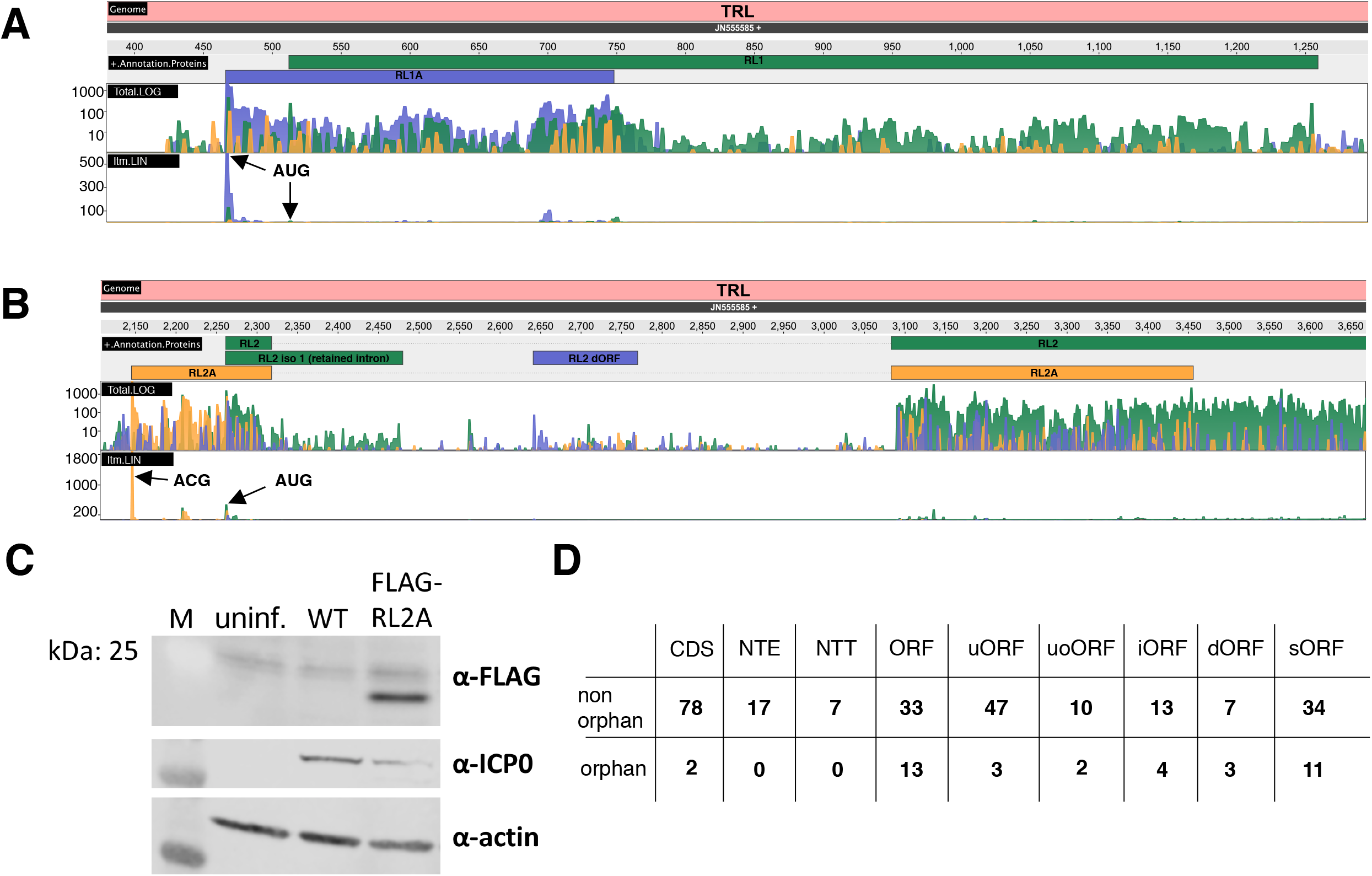
Ribosome profiling data visualizing expression of two novel viral ORFs (RL1A, RL2A) expressed from the **(A)** RL1 and **(B)** RL2 locus of the terminal repeats (TRL). Both standard ribosome profiling data in log scale (Total.log) as well as translation start site profiling data obtained using Lactimidomycin (linear scale, ltm.LIN) are shown. Both ORFs are well expressed and validated by the strong peak of their respective TaSS in the ltm-track (black arrows). While RL2A initiates from a non-canonical ACG start codon, the 93 aa RL2A protein initiates from an AUG start codon and was previously missed due to its length of <100 aa. **(C)** Validation of RL2A expression by Western blot. Primary human fibroblasts were infected with virus inoculum produced from transfection of bacterial artificial chromosomes (BACs) encoding either wild-type HSV-1 (WT) or a virus expressing a FLAG-tagged-RL2A (FLAG-RL2A). Western blot analysis was performed at 24 h post infection. Expression of ICP0 and actin served as infection and house-keeping control, respectively. **(D)** Distributions of all identified types of ORFs of HSV-1 classified by ORFs and orphan ORFs.

A second interesting novel ORF is expressed in the RL2 locus, which encodes the major viral immediate early protein lCP0. Here, we identified a novel spliced ORF (termed RL2A) of 181 aa that initiates from an ACG start codon 116 nt upstream of the lCP0 TaSS (Fig. 6b). Expression of RL2A was confirmed by generating a mutant virus with a triple-FLAG-tag inserted 12 nt downstream of the ACG start codon (Fig. 6c). lnterestingly, RL2A expression by the mutant virus could only be detected immediately upon virus reconstitution and was readily lost upon serial passaging indicating that insertion of the triple-FLAG-tag into the RL2A repeat region severely impaired viral fitness.

Transcription of all viral genes continuously increases throughout lytic infection with the exception of the transcript encoding ORF-O and ORF-P. These two partially overlapping ORFs are expressed antisense to the lCP34.5 (RL1) gene^44^. Consistent with the previous report, the respective transcript was already well detectable in 4sU-seq data at 1 h p.i. but transcriptional activity declined rapidly afterwards (Suppl. Fig. 7a). Nevertheless, translation of the respective transcripts remained detectable until late times of infection. lnterestingly, the absence of a canonical start codon resulted in the hypothesis that ORF-O initiates from the same AUG start codon as ORF-P but then diverges within the first 35 codons due to a ribosomal frame shift. We did not observe any evidence for frame-shifts in the HSV-1 translatome. While translation of ORF-O was only rather weak, our data indicate that it rather initiates from a non-canonical ACG start codon 76 nt upstream of the ORF-P (Suppl. Fig. 7b).

Finally, we also aimed to validate novel HSV-1 ORFs by whole proteome mass spectrometry. We obtained triple-SlLAC whole proteome data from HSV-1 infected HFF at 0, 4 and 8 h p.i (n=3). Furthermore, we performed whole proteome mass spectrometry from primary lung fibroblasts infected with HSV-1 for 0, 4, 9 and 15 h. ln total, this confirmed only 10 (5.5%) of the 186 novel ORFs and sORFs (Suppl. Tab. 3c; excluding NTEs and NTTs). This rather small fraction is consistent with previous work on novel HCMV ORFs^2^ and, presumably, reflects that the majority of viral sORFs are inherently unstable and rapidly degraded upon translation similar to their cellular counterparts. Nevertheless, they may play an important role in regulating translation of the viral ORFs encoded downstream of them. Furthermore, the novel large viral ORFs were expressed at substantially lower (~10x) levels than the previously identified viral protein coding sequences (Suppl. Fig. 8).

### Development of a new integrative nomenclature of HSV-1 gene products

The large number of novel viral gene products required the extension of the current nomenclature. We annotated the novel viral gene products and compiled viral gene units comprising transcript isoforms, ORFs and regulatory entities, e.g. uORFs and uoORFs (Suppl. Tab. 3a). A detailed description of the applied rules is provided in Suppl. Methods. In brief, we fully maintained the current nomenclature for all ORFs in the reference annotation^1^ and attributed each ORF to the most highly expressed transcript in its vicinity. Alternative transcript isoforms initiating within less than 500 nucleotides were labeled with “*” (extended) or “#” (truncated), e.g. UL13 mRNA *1. Finally, alternatively spliced transcripts were labeled with “iso 1” and “iso 2”. Short ORFs (<100aa), were named upstream ORF (uORF), upstream overlapping ORF (uoORF), internal ORF (iORF) and downstream ORF (dORF) in relation to the next neighboring large ORF. Any ORF for which no transcript could be identified to be responsible for its translation was labeled as “orphan”. An overview of the status of the various kinds of ORFs we identified is shown in Fig. 6d. In accordance, any transcript, which was not found to encode an ORF or sORF within its first 500 nt was also labeled as “orphan”. Interestingly, we identified 41 “orphan” transcripts (Suppl. Tab. 6), which showed predominantly nuclear localization indicating that they may represent novel viral nuclear long non-coding RNAs (lncRNAs). However, all of them were expressed at rather low levels. Accordingly, we were unable to validate five of them by Northern blots despite extensive efforts. We conclude that HSV-1 does not express any highly transcribed viral non-coding RNAs during lytic infection.

## Discussion

In recent years, major advances in high-throughput experimental methodology have revealed that herpesvirus gene expression is surprisingly complex. While a number of studies in the last few years described hundreds of novel viral transcripts and ORFs, a systematic analysis, validation and integration into gene modules, which attribute individual ORFs and sORFs to specific transcripts they are expressed from, was not attempted. Moreover, the lack of a standardized nomenclature has hampered functional studies on these new viral gene products. Based on a wide spectrum of new and published functional genomics data, we here provide a state-of-the-art, fully revised annotation of the HSV-1 genome.

Calling novel gene products based on big data poses the risk of false positives. Integrative analysis of transcription start site (TiSS) data obtained by both second (cRNA-seq, dRNA-seq) and third (PacBio, MinION) generation sequencing approaches highlighted the necessity to validate viral TiSS by multiple means to exclude such experimental artifacts generated by the individual approaches. Similarly, the different transcription profiling approaches identified numerous putative novel splicing events. However, the vast majority of these only occurred at very low levels. We restricted our analysis to the first 8 h of lytic HSV-1 infection and only included splicing events observed by at least two approaches. While MinION sequencing recently identified novel intergenic splice sites resulting in novel fusion proteins (e.g. between ICP0 and glycoprotein L)^29^, the respective transcripts only arise very late in infection and their functional relevance remains unclear. We thus did not include them into our annotation. We conclude that splicing in the HSV-1 transcriptome was already well described by the previous reference annotation but rare splicing events may explain some of our “orphan” viral ORFs and sORFs.

In eukaryotic cells, RNA polymerase II (Pol II) may continue transcribing for thousands of nucleotides downstream of the PAS until transcription is terminated and Pol II is released from the chromatin^45^. With viral gene expression rapidly increasing throughout lytic infection, mRNA precursors with unprocessed 3’-ends that still extend beyond the canonical polyadenylation site are likely to be prevalent in the infected cells. Thus, unprocessed viral pre-mRNAs can easily be misinterpreted as mature viral transcripts. They presumably explain previous reports of near-complete transcription of herpesviral genomes during productive infections^2,46,47^. Analysis of cytoplasmic rather than total RNA provides a more accurate picture of the mature viral transcriptome. Consistent with previous reports, we confirmed that the major viral RNA export factor ICP27 was required for efficient export of all viral transcripts^13^. Interestingly, this also included all immediate early genes and spliced viral transcripts.

Ribosome profiling identified 134 novel sORFs expressed during lytic HSV-1 infection. The vast majority of these represent so called upstream open reading frames (uORFs). Interestingly, a relatively large fraction of transcript isoforms (~20%) encode their own uORF, which preferentially (54%) initiate from AUG start codons. Cellular uORFs constitute an important regulatory network governing gene expression at the level of translation by affecting translation initiation of the downstream ORF^48^. Reliable annotation of both viral transcripts and their respective uORFs will now enable functional studies on these cryptic viral gene products. sORF-encoded polypeptides are usually highly unstable and thus remain undetectable by whole proteome mass spectrometry (WP-MS). Accordingly, we were only able to confirm about 5.5% of our novel ORFs by WP-MS. Interestingly, we could recently show that peptides derived from cellular sORFs are nevertheless efficiently incorporated into and presented by MHC-I molecules on the cell surface despite remaining virtually undetectable by WP-MS^5^. sORF-derived peptides thus may constitute a new viral class of antigens that are efficiently presented by MHC-I but, due to their instability and extremely low abundance within the cell’s proteome, will represent poor substrates for cross-presentation and CD4-CD8 augmentation. Further studies are necessary to assess the role of HSV-1 sORFs in the regulation of viral protein expression, antiviral T cell control and evasion thereof.

Based on our revised annotation of 201 viral transcripts and 284 ORFs, we extended the existing nomenclature to include these novel viral gene products. This did not involve any renaming of previously described viral gene products. Our nomenclature thereby explains gene expression of the majority of viral ORFs in the context of different transcript isoforms, uORF and uoORFs. This will facilitate functional studies on the novel viral gene products as well as their transcriptional and translational regulation.

## Methods

### Cell culture, viruses and infections

Human foreskin fibroblasts (HFF, #86031405, purchased from ECACC), 293T, Vero 2-2 (Smith, Hardwicke, & Sandri-Goldin, 1992) BHK-21 and BHK-21 dox-UL19 (described below) cell lines were cultured in flasks containing Dulbecco’s Modified Eagle Medium (DMEM), high glucose, pyruvate (ThermoFisher #41966052) supplemented with 1x MEM Non-Essential Amino Acids (ThermoFisher #11140050), 1mM additional sodium pyruvate (ThermoFisher #11360070), 10% (v/v) Fetal Bovine Serum (FBS, Biochrom #S 0115), 200lU/mL, penicillin (pen) and 200μg/mL streptomycin (strep). All cells were incubated at 37°C in a 5% (v/v) CO_2_-enriched incubator.

HFF were utilized from passage 11 to 17 for all high-throughput experiments. Virus stocks for wild-type HSV-1 were produced on baby hamster kidney (BHK) cells as described. Stocks of the ICP27 null mutant (strain KOS)^49^ were produced on complementing Vero 2-2 cells^50^. HFF were infected for 15 min at 37°C about 24 h after the last split using a multiplicity of infection (MOl) of 10. Subsequently, the inoculum was removed and fresh media was applied to the cells.

To reconstitute the FLAG-tagged UL19 NTE, BHK-21 dox-UL19 cells with doxycycline-inducible expression of UL19 were generated by cloning the HSV-1 Syn17+ UL19 coding sequence using primers described in Suppl. Tab. 7 into the Sall and Nhel sites of pTH3, a derivative of pCW57.1 with a custom multiple cloning site in lieu of the gateway cloning site and the addition of the TRE tight promoter from pTRE-Tight. Lentiviral vectors were generated by cotransfection of this construct with psPAX2 and pCMV-VSV-G into 293T cells. Lentivirus-containing supernatants were sterile-filtered with Minisart^®^ NML 0.45μm cellulose acetate filters (Sartorius #17598) and added to BHK-21 cells. Polyclonal populations were selected 48 h post-transduction and maintained in 1μg/mL puromycin.

### Viral mutagenesis and reconstitution

All viral mutants were generated via *en passant* mutagenesis^51^ using *Escherichia coli* strain GS1783 with the bacterial artificial chromosome (BAC) HSV1(17+)-LoxCheVP26^52^ expressing a fusion protein of mCherry on the N-terminus of the UL35 gene product (VP26). Full primer and construct sequences can be found in the Suppl. Tab. 7. BAC DNA was purified using the NucleoBond BAC 100 kit (Macherey-Nagel #740579) and transfected for virus reconstitution into BHK-21 cells with Lipofectamine 3000 (ThermoFisher #L3000-075). HSV-1 expressing the FLAG-tagged N-terminal extension of UL54 virus were reconstituted and titrated on Vero 2-2 cells^50^. The virus expressing the tagged N-terminal extension of UL19 was generated in BHK-21 dox-UL19 cells. BHK-21 dox-UL19 cells were plated the day before in media containing 1μg/mL doxycycline (Sigma #D3072), which was maintained throughout virus generation.

Virus produced by transfected cells was expanded on minimally five T175 flasks of the corresponding cell type. Virus-containing supernatants were harvested upon >90% cytopathic effect and centrifugation at 8,000 RCF at 4°C for 10 min to pellet cells. Cell pellets were snap-frozen in liquid nitrogen and thawed at 37°C three times to free cell-associated virus. Cellular debris was pelleted at 10,000 RCF, 4°C for 10 min and supernatant combined with the supernatant in the previous step. Virions were pelleted by centrifugation at 19,000 RCF for two hours at 4°C, resuspended in phosphate-buffed saline (PBS), and pelleted once more over a 20% (w/v) sucrose cushion in PBS 16,000 RPM for two hours at 4°C in a SW 28 swinging-bucket rotor (Beckman). Virus pellets were resuspended in PBS, snap-frozen in liquid nitrogen, stored at −80°C, and titrated by plaque assay. Infections were carried out in serum-free DMEM containing penicillin and streptomycin for 1 h at 37°C. The time at which inoculum was replaced with growth media was marked as the 0 h timepoint.

### Western blot

Samples were harvested at the indicated timepoints by removal of growth media and direct lysis in 2x Laemmli buffer containing 5% (v/v) β-mercaptoethanol. Samples were sonicated and heated for 5 min at 95°C before loading onto a Novex™ WedgeWell™ 4-20% Tris-Glycine Gel (ThermoFisher #XP04200BOX). Proteins were transferred to polyvinylidene difluoride (PVDF) membranes, blocked for 1 h at room temperature in 1xPBST containing 5% (w/v) milk (Carl Roth T145.3), and probed using α-FLAG M2 (Sigma #F1804) overnight at 4°C at a 1:1000 at dilution and α-mouse IgG (whole molecule)-peroxidase (Sigma #9044) for 1 h. β-actin was probed using α-β-actin C4 antibody (Santa Cruz #sc-47778) at a 1:1000 dilution for 1 h, followed by IRDye^®^ 800CW goat α-mouse IgG (LI-COR #926-32210) at 1:5000 for 1 h. ICP0 was probed using α-ICP0 clone 5H7 (Santa Cruz #sc-56985) at a 1:1000 dilution for 1 h, followed by lRDye^®^ 680RD goat α-mouse lgG (Ll-COR #926-68070) at 1:5000 for 1 h. Samples were washed with 1xPBST and blocked before addition of each antibody in the milk/PBST buffer. Blots were visualized with a Ll-COR Odyssey^®^ FC lmaging System.

### Immunofluorescence

10^5^ HFF cells were plated on glass coverslips in 12-well dishes 24h prior to infection. At 8 h post infection cells were fixed in 4% formaldehyde in PBS for 1 h at room temperature, washed three times in PBS and stored at 4°C overnight in PBS. Cells were incubated in permeabilization buffer (10% FBS, 0.25M glycine, 0.2% Triton X-100, 1xPBS) for 1 h at room temperature before incubating them in blocking buffer (10% FBS, 0.25M glycine, 1xPBS) for 1 h at room temperature. Anti-FLAG antibody (GenScript #A00187) was incubated in 10% FBS and 1xPBS for 1 h at 37°C at a concentration of 1μg/mL. The secondary anti-mouse lgG, Alexa Fluor 488 (ThermoFisher #A11017) was incubated in 10% FBS in 1xPBS for 1 h at room temperature with 0.5μg/mL 4’,6-diamidino-2-phenylindole (DAPl). All steps were followed by three 5-minute washes in PBS except for after the primary antibody, which was washed with 1xPBS and 0.05% Tween-20. Coverslips were washed in water before mounting them in medium containing Mowiol 4-88 and 2.5% (w/v) 1,4-diazabicyclo[2.2.2]octane (DABCO).

### Transcription start site (TiSS) profiling

Total cellular RNA was isolated using Trizol (lnvitrogen) following the manufacturer’s instructions. RNA was resuspended in water and stored at −80°C until use. TiSS profiling using cRNA-seq was performed as described for HCMV^2^. TiSS using dRNA-seq was performed according to the published protocol^10^ with some modifications by the Core Unit Systems Medicine (Würzburg). ln brief, for each sample 3 μg of DNase-digested RNA was treated with T4 Polynucleotide Kinase (NEB) for 1 h at 37 °C. RNA was purified with Oligo Clean & Concentrator columns (Zymo) and each sample was split into an Xrn1 (+Xrn1) and a mock (-Xrn1) sample. The samples were treated with 1.5 U Xrn1 (NEB; +Xrn1) or water (-Xrn1) for 1 h at 37 °C. Digest efficiency was checked on a 2100 Bioanalyzer (Agilent) and 5’ caps were removed by incubation with 20 U of RppH (NEB) for 1 h at 37 °C. Afterwards RNA was purified and eluted in 7 μl. 6 μl were used as input material for the NEBNext^®^ Multiplex Small RNA Library Prep Set for lllumina^®^. Library preparation was performed according to the manufacturer’s instruction with the following modifications: 3’ adapter, SR RT primer and 5’ adapter were diluted 1:2, 13 cycles of PCR were performed with 30 sec of elongation time, and no size selection was performed at the end of library preparation. Concentrations of libraries were determined using the Qubit 3.0 (Thermo Scientific) and their fragment sizes were determined using the Bioanalyzer. Libraries were pooled equimolar. Sequencing of 75 bp single-end reads was performed on a NextSeq 500 (Illumina) at the Cambridge Genomic Services (cRNA-seq) and the Core Unit Systems Medicine in Würzburg (dRNA-seq). To validate TiSS identified by cRNA-seq, dRNA-seq, PacBio or MinION, total RNA-seq and 4sU-seq data that were previously published^9^ were reanalyzed (see below).

### RNA-seq of subcellular RNA fractions

Subcellular RNA fractions (cytoplasmic, nucleoplasmic and chromatin-associated RNA) were prepared by combining two previously published protocols^53,54^. Data from uninfected and wild-type HSV-1 infected cells were published recently^30^. Infection with the ICP27-null mutant was performed in the same experiment. As for wild-type HSV-1 infection, total cellular RNA was isolated using Trizol at 8 h p.i. The efficiency of the fractionations was controlled by qRT-PCRs for intron-exon junctions for ACTG1 (chromatin-associated vs the other three fractions) and western blots for histone H3 (nuclear vs cytoplasmic fraction) (data not shown). Fractionation efficiencies were furthermore confirmed on the RNA-seq data by comparing expression values of known nuclear and cytoplasmic RNAs as well as intron contributions^30^. Sequencing libraries were prepared using the TruSeq Stranded Total RNA kit (Illumina) following rRNA depletion using Ribo-zero. Sequencing of 75 bp paired-end reads was performed on a NextSeq 500 (Illumina) at the Cambridge Genomic Services and the Core Unit Systems Medicine (Würzburg).

### Ribosome profiling

The ribosome profiling time-course data (lysis in presence of cycloheximide) have already been published^55^. Additionally, so far unpublished data we generated include translation start site (TaSS) profiling performed by culturing cells in medium containing either Harringtonine (2 μg/ml) or Lactimidomycin (50 μM) for 30 min prior to harvest. Harringtonine samples were obtained for 2 h and 8 h p.i., Lactimidomycin was employed for mock, 4 and 8 h p.i. Two replicates of each condition were analyzed. All libraries were sequenced on a HiSeq 2000 at the Beijing Genomics Institute in Hong Kong.

### Proteomic analysis

#### Stable Isotope Labelling with Amino Acids in Cell Culture (SILAC)

Primary human foreskin fibroblasts were grown for 5 passages in DMEM lacking lysine and arginine (Thermo Scientific) supplemented with 10 % dialysed FCS (Gibco), 100 units/ml penicillin and 0.1 mg/ml streptomycin, 280 mg/L proline (Sigma) and light (K0, R0; Sigma), medium (K4, R6; Cambridge Isotope Laboratories) or heavy (K8, R10; Cambridge Isotope Laboratories) 13C/15N-containing lysine (K) and arginine (R) at 50 mg/L.

#### Infection with HSV-1

Pre-labeled cells were infected with HSV-1 at a multiplicity of infection (MOI) of 10 for 4 or 8 hrs, and uninfected cells were included as a control. The experiment was conducted in triplicate (biological replicates), with a 3-way SILAC label swap.

#### Sample preparation

Cells were harvested at the indicated time points, washed with ice-cold PBS, snap-frozen in liquid nitrogen and stored at −80 °C prior to filter-aided sample preparation (FASP) essentially as previously described^56,57^. In brief, cell pellets were lysed in 4% SDS/100 mM Tris HCl pH 7.4 supplemented with cOmplete protease inhibitor cocktail (Roche), and sonicated at 4 °C using a Diagenode Bioruptor. Protein concentrations were determined using the Pierce BCA Protein Assay kit (Thermo Scientific) and light, medium and heavy lysates combined 1:1:1 by protein mass (total 45 μg protein/replicate). Lysates were transferred to Microcon-30 kDa centrifugal filter units (Millipore), reduced (100 mM DTT) and alkylated (50 mM iodoacetamide) at room temperature, washed with a total of 5 column volumes of 8 M urea/100 mM Tris HCl pH 8.5, then digested with 1 μg modified sequencing grade trypsin (Promega) in 50 mM ammonimum bicarbonate at 37 °C for 12 hrs. Peptide eluates were collected by centrifugation and stored at −80 °C prior to fractionation.

#### Off-line High pH Reversed-Phase (HpRP) peptide fractionation

HpRP was conducted using a Dionex UltiMate 3000 UHPLC system (Thermo Scientific) powered by an ICS-3000 SP pump with an Agilent ZORBAX Extend-C18 column (4.6 mm × 250 mm, 5 μm particle size). Peptides were resolved using a linear 40 min 0.1 %-40 % MeCN gradient at pH 10.5 with eluting peptides collected in 15 s fractions. Peptide-rich fractions were concatenated across the gradient to give 10 pooled fractions/sample, dried using an Eppendorf Concentrator then re-suspended in 15 μL mass spectrometry solvent (3 % MeCN, 0.1 % TFA).

#### Mass spectrometry

Data were generated using an Orbitrap Fusion Tribrid mass spectrometer (Thermo Scientific). Peptides were fractionated using an RSLCnano 3000 (Thermo Scientific) with solvent A comprising 0.1% formic acid and solvent B comprising 80% MeCN, 20% H_2_O, 0. 1% formic acid. Peptides were loaded onto a 50 cm Acclaim PepMap C18 column (Thermo Scientific) and eluted using a gradient rising from 3 to 40% solvent B by 73 min at a flow rate of 250 nl/min. MS data were acquired in the Orbitrap at 120,000 fwhm between 350–1500 m/z. Spectra were acquired in profile with AGC 5 × 10^5^. Ions with a charge state between 2+ and 7+ were isolated for fragmentation in top speed mode using the quadrupole with a 1.6 m/z isolation window. HCD fragmentation was performed at 33% collision energy with fragments detected in the ion trap between 350–1400 m/z. AGC was set to 5 × 10^3^ and MS2 spectra were acquired in centroid mode.

### Data analysis, statistics and reproducibility

Random and sample barcodes in cRNA-seq and ribosome profiling data were analyzed as described^9^. Briefly, barcodes and 3’ adapters were trimmed from the reads using our inhouse computational genomics framework gedi (available at https://github.com/erhard-lab/gedi). Reads were mapped using bowtie 1.0 against the human genome (hg19), the human transcriptome (Ensembl 75) and HSV-1 (JN555585). All alignments for a read with minimal number of mismatches were filtered and assigned to specific samples based on the sample barcode. PCR duplicates were collapsed based on random barcodes while paying attention to sequencing errors.

dRNA-seq data was processed in a similar manner to the cRNA-seq and ribosome profiling data with the exception that PCR duplicates were not collapsed as no random barcodes were used. 4sU-seq, total RNA-seq and RNA-seq data of subcellular fractions were analyzed as described^30^

TiSS profiling data were analyzed with the newly developed tool iTiSS (<u>i</u>ntegrative <u>T</u>ranscr<u>i</u>ptional <u>S</u>tart <u>S</u>ite caller; manuscript in preparation). Briefly, dRNA-seq and cRNA-seq data were searched for positions showing strong accumulation of reads 5’ ends compared to their surroundings using a sliding-window approach. Transcriptional activity was defined by having a stronger transcriptional activity downstream of a putative TiSS than upstream identified using Fisher’s exact test. Selective induction or repression (altered expression kinetics) of the respective transcripts during the course of HSV-1 infection was defined by observing a significant change of transcriptional activity surrounding a putative TiSS (100 bp window) during the course of infection using a likelihood ratio test based on the dirichlet distribution.

The 9 different points a single TiSS can score are defined as follows: (1) A significant accumulation of reads in both replicates of the cRNA-seq dataset, (2) the presence of a stronger transcriptional activity downstream than upstream in both replicates, (3) the presence of a significant change during the course of infection in both replicates, (4) a significant accumulation of reads in the dRNA-seq dataset in both replicates, (5) a TiSS called in the PacBio data set in close proximity (+/− 5 nt). For this purpose, we manually corrected the GFF-file provided alongside the GEO-submission for the PacBio data, which was inconsistent with the transcripts reported in the paper. (6) A TiSS called in the MinlON data set with a maximum distance of 20 nt downstream, (7) the presence of an ORF at most 250 bp downstream, which is not yet explained by another transcript, (8) significantly different expression kinetics throughout the viral infection in the 4sU dataset and (9) a significant increase in read coverage downstream in the 4sU dataset.

The reported enrichment factors for dRNA-seq and cRNA-seq were calculated based on predicted TiSS in human rather than HSV-1. This was done to prevent undesired biases due to read-in caused by the extraordinary high number of overlapping transcripts in HSV-1. The predicted TiSS were ordered based on the number of reads starting at their respective positions. The median was then calculated over the 50 strongest and 10 strongest expressed TiSS for cRNA-seq and dRNA-seq, respectively.

Significance of the correlation between the presence of a TATA-box-like motif and the transcription strength of TiSS was calculated using Fisher’s exact test. Here, the TiSS were ordered by their numbers of reads starting and sorted into three equally sized bins. For the bin containing the strongest TiSS as well as the bin containing the weakest TiSS, the number of all nucleotides between position −30 and −25 relative to the TiSS were summed up. For the parameters of the Fisher’s exact test, the following sums were used a=T+A(strongest bin), b=C+G(strongest bin), c=T+A(weakest bin) and d=C+G(weakest bin).

We used our in-house tool PRICE^5^ version 1.0.1 to call ORFs separately for the two replicates of ribosome profiling data but pooling all samples from each replicate.

RNA-seq data were mapped using STAR^58^ version 2.5.3a using a combined reference index derived from Ensembl 90 and our final HSV-1 annotation.

We analyzed mass spectrometry data using MaxQuant^59^ version 1.6.5.0. Spectra were matched against a combined database of proteins from Ensembl (version 75), and all ORFs identified by ribosome profiling. We used carbamidomethylation as fixed and acetylation (N-terminal) and oxidation at methionine as variable modifications. Peptides were filtered for 1% FDR using the target-decoy approach by MaxQuant.

The export indices of chromatin-associated RNA and cytoplasmic RNA were derived by computing their fold changes between the wild type and the null mutant for ICP27 using the lfc R-package^60^.

Analysis of the predicted protein sequences to reveal function and functional motifs used sequence comparison, domain composition, structure prediction and motif searches as described previously^61–63^.

## Supporting information

Principles of new nomenclature

## Reporting Summary

Further information on research design is available in the Nature Research Reporting Summary linked to this article.

## Code availability

Scripts used to create the Figures and Tables and analyze the omics data are available at zenodo (doi: 10.5281/zenodo.2621226).

## Data availability

All sequencing data have been deposited in the Gene Expression Omnibus (GEO) database with accession codes GSE128324 (Translation start site profiling, transcription start site profiling), GSE59717 (4sU-seq and total RNA-seq), GSE60040 (ribosome profiling) and GSE128880 (cytoplasmic, nucleoplasmic and chromatin-associated RNA). Additional data from other sources utilized in this work are available at the GEO database with accession code GSE97785 (PacBio)^11^, and from the European Nucleotide Archive (ENA) with the study accession code PRJEB27861 (MinION)^12^. The mass spectrometry proteomics data have been deposited to the ProteomeXchange Consortium via the PRIDE [1] partner repository with the dataset identifiers PXD013010 and PXD013407.

## Acknowledgments

This work was supported by the European Research Council (ERC-2016-CoG 721016 – HERPES to L.D.), the MRC (CSF G1002523 to L.D. and MR/P008801/1 to N.J.M.), the Wellcome Trust (PRF 210688/Z/18/Z to P.J.L.), NHSBT (WP11-05 to L.D. and WPA15-02 to N.J.M.), DFG (1275/6-1 to L.D., GR950/16-1 to F.G., LA2941/4-1 to M.L., SFB1123/Z2 to R.Z. and FR2938/1-2 to C.C.F.), the NIHR Cambridge BRC, a Wellcome Trust Strategic Award to CIMR and the IZFK at the University of Würzburg (project Z-6). A.W.W. was the recipient of a generous grant from the Alexander von Humboldt Foundation and the German Federal Foreign Office.

## Authors Contributions

A.W.W., T.H., E.W., B.P., A.L.H., M.G., K.D., J.M., A.J.R., R.A., N.M., F.W.H.K, G.M., C.B., S.K., and F.G. performed the experiments. A.W.W., C.S.J., F.E. and L.D. designed the experiments, analyzed the data and wrote the paper. C.S.J., C.C.F. and F.E. performed the computational analyses. R.Z. supervised development of the computational methods (for ribosome profiling analysis). S.K., M.L. and P.J.L. supervised the mass spectrometry analyses and L.C. and T.D. contributed to motif analysis for the novel viral ORFs.

## Corresponding author

Correspondence to Lars Dölken and Florian Erhard.

## Supplementary information

Supplementary Tables 1 – 7, Supplementary Figures 1 – 8.

Reporting Summary

## Supplementary Figures

**Supplementary Figure 1. Comparison of transcription start sites obtained by PacBio and MinION**

**(A)** Distance of transcription start sites (TiSS) identified by PacBio^11^ to the TiSS positions obtained by both cRNA-seq and dRNA-seq is shown in relation to read counts. This confirmed TiSS identified by cRNA-seq and dRNA-seq at single nucleotide resolution. (B) Same as for **(A)** but for TiSS obtained from MinlON sequencing data^12^. The 98 TiSS called by MinlON generally lacked 7-18 nucleotides (nt) at the 5’ end for technical limitations of the MinlON direct RNA sequencing method. After correcting for this, the manually curated MinlON data confirmed many of the novel TiSS we identified.

**Supplementary Figure 2. Identification of splicing events in the HSV-1 transcriptome**

Shown are the locations of known and putative splicing events in the HSV-1 transcriptome depicted by their unique read counts for the upstream, exon-spanning and downstream sequences. Besides the 8 known splicing events and a novel NAGNAG event in the lCP0 mRNA, lllumina sequencing confirmed 4 splicing events identified by PacBio^11^. Furthermore, screening our RNA-seq and 4sU-seq data for novel splicing events with at least 10 nt exon-spanning reads identified 58 putative additional splicing events. However, exon-spanning reads were >10-fold less prevalent than reads mapping to the flanking regions. We therefore, decided not to include them into our new reference annotation. Nevertheless, the may explain some of the “orphan” ORFs. Our data confirmed all 11 splicing events. However, only four of them occurred at relevant levels and were included into our new reference annotation.

**Supplementary Figure 3: Read-through of the UL24 polyadenylation site**

Screenshot of our HSV-1 viewer depicting the read-through of the UL24 polyadenylation site. This is already visible at 2 h p.i. and is used express the UL25 RNA transcript, which translates for the UL25 CDS.

**Supplementary Figure 4: Large truncated viral ORFs**

Screenshots of our HSV-1 viewer depicting the seven viral ORFs with NTEs. Alternative TiSS downstream of the main TiSS explained translation of 6 of 7 NTTs. Only for US3.5, we could not identify a corresponding transcript. The NTTs as well the original ORFs are highlighted in pink. The corresponding transcripts for both are highlighted in red. Red arrows in the cRNA-seq and dRNA-seq track point at the specific TiSS validation, if present. Pink arrows point at the TaSS validation if present. **(A)** NTTs encoded in the sense strand. **(B)** NTTs encoded in the antisense strand. For UL29/UL29.5 the combined ribosome profiling data for 1 h p.i. up to 8 h p.i. is shown instead of the Lactimidomycin. This was a pure esthetical decision as the TaSS peak there was too prominent and therefore the translation throughout the ORF could not be seen anymore.

**Supplementary Figure 5: Validation of N-terminal extensions by IF**

Immunofluorescence (IF) of human foreskin fibroblasts infected with Vp26-mCherry HSV-1 containing FLAG-tags inserted upstream of the canonical start codon into the N-terminal extension (NTE) or downstream of it (AUG) for US5 **(A)** and UL54 **(B)**. Cell nuclei where stained using DAPI. No differences in the subcellular localization between the NTE and the canonical protein were observed.

**Supplementary Figure 6: Prediction of alphaherpesviral US3 N-terminal extensions.**

Primary peptide sequences for validated (HSV-1, PRV) and predicted (HSV-2, BHV-1, FeHV-1 and MaHV-1) US3 NTEs are depicted from the start codons (canonical or non-canonical) to the annotated US3 start codon (“M” in bold). Hydrophobic resides are indicated in red. Putative nuclear export signals matching the motif [LIVFM]-X_2,3_-[LIVFM]-X_2,3_-[LIVFM]-X-[LIVFM] are highlighted in yellow. The mitochondrial localization signal predicted for PRV^39^ is underlined.

**Supplementary Figure 7. Expression of ORF-O and ORF-P**

**(A)** Expression kinetics of the ORF-O/ORF-P mRNA depicted by cRNA-seq. While the mature transcript is well expressed at 1 h p.i. Transcriptional activity rapidly declines afterwards and is obscured by transcription from upstream genomic regions later on in infection. **(B)** Ribosome profiling data for ORF-O and ORF-P. Combined data from all time points analyzed by standard ribosome profiling (Total log) is shown to account for the overall low translation rates. While translation of ORF-P is well represented, translation of ORF-O is less prominent. While we cannot fully exclude the previously proposed frameshift within ORF-O, a strong translation start site peak obtained by Lactimidomycin treatment 76 nt upstream of the AUG start codon of ORF-P is consistent with ORF-O initiating from an ACG start codon upstream of ORF-P.

**Supplementary Figure 8: Expression strength of all identified large (≥100 aa) ORFs**

Expressions strengths of all ORFs with an amino acid length ≥100 over the course of the infection classified by their respective ORF type. This includes all known large ORFs (CDS) and 41 ORFs and 2 iORFs. Overlapping ORFs translated in the same frame were excluded. All newly identified ORFs show lower expressions than the previously identified ones (CDS). Most of the novel large ORFs are expressed at relatively low levels compared to the known large viral ORFs.

**Supplementary Tables**

**Supplementary Table 1. HSV-1 transcripts**

**Supplementary Table 2. HSV-1 splicing events**

**Supplementary Table 3. HSV-1 ORFs / Predicted function / MS validation**

**Supplementary Table 4. Truncated HSV-1 ORFs**

**Supplementary Table 5. HSV-1 ORFs with N-terminal extension (NTE)**

**Supplementary Table 6. HSV-1 orphan ORFs**

**Supplementary Table 7. Primers and gene synthesis constructs**

