## Supplementary Figures for "Integrative functional genomics decodes herpes simplex virus 1"

A

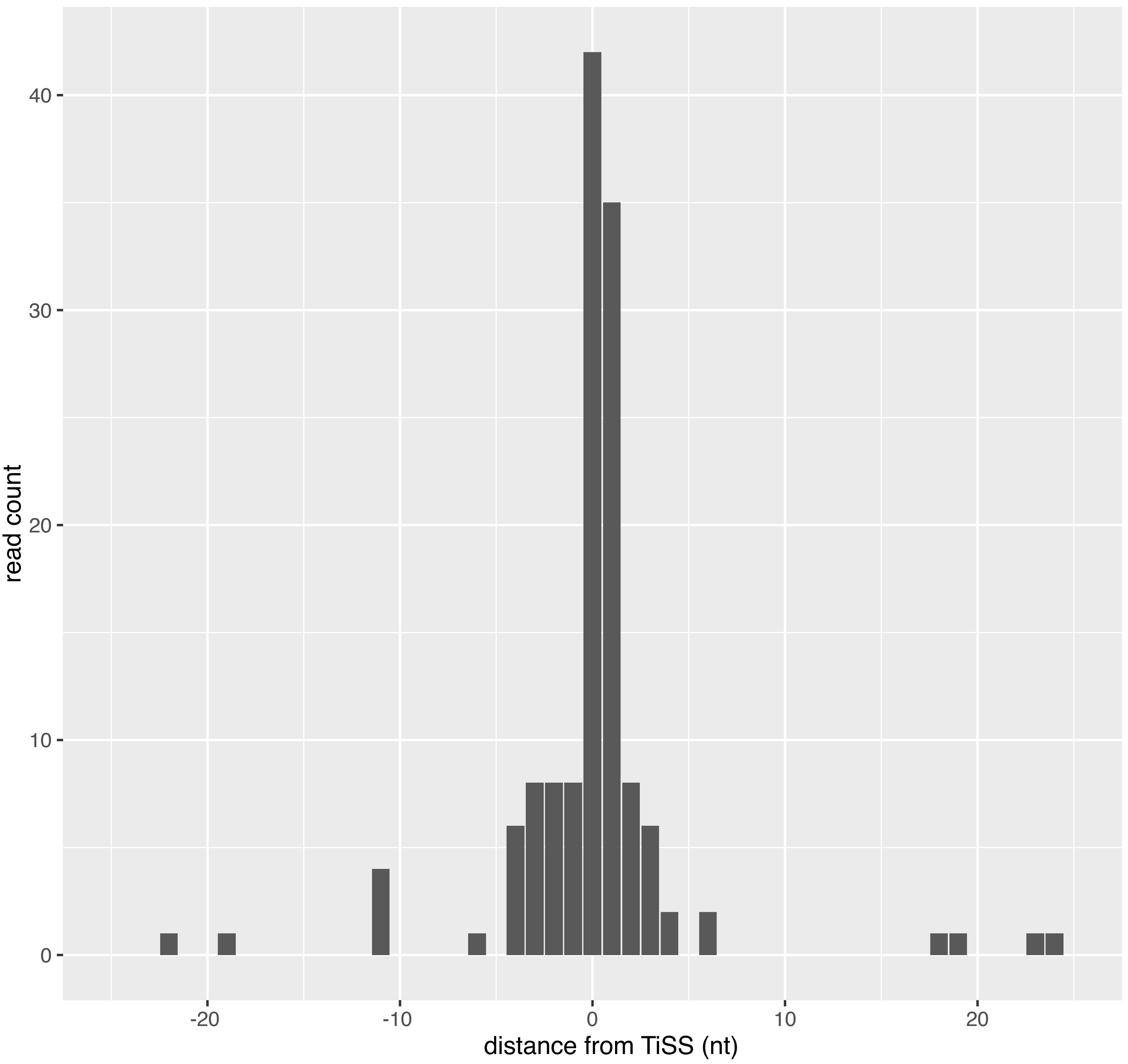

40
30
20
10
0
read count
-20
-10
0
10
20
distance from TiSS (nt)


B

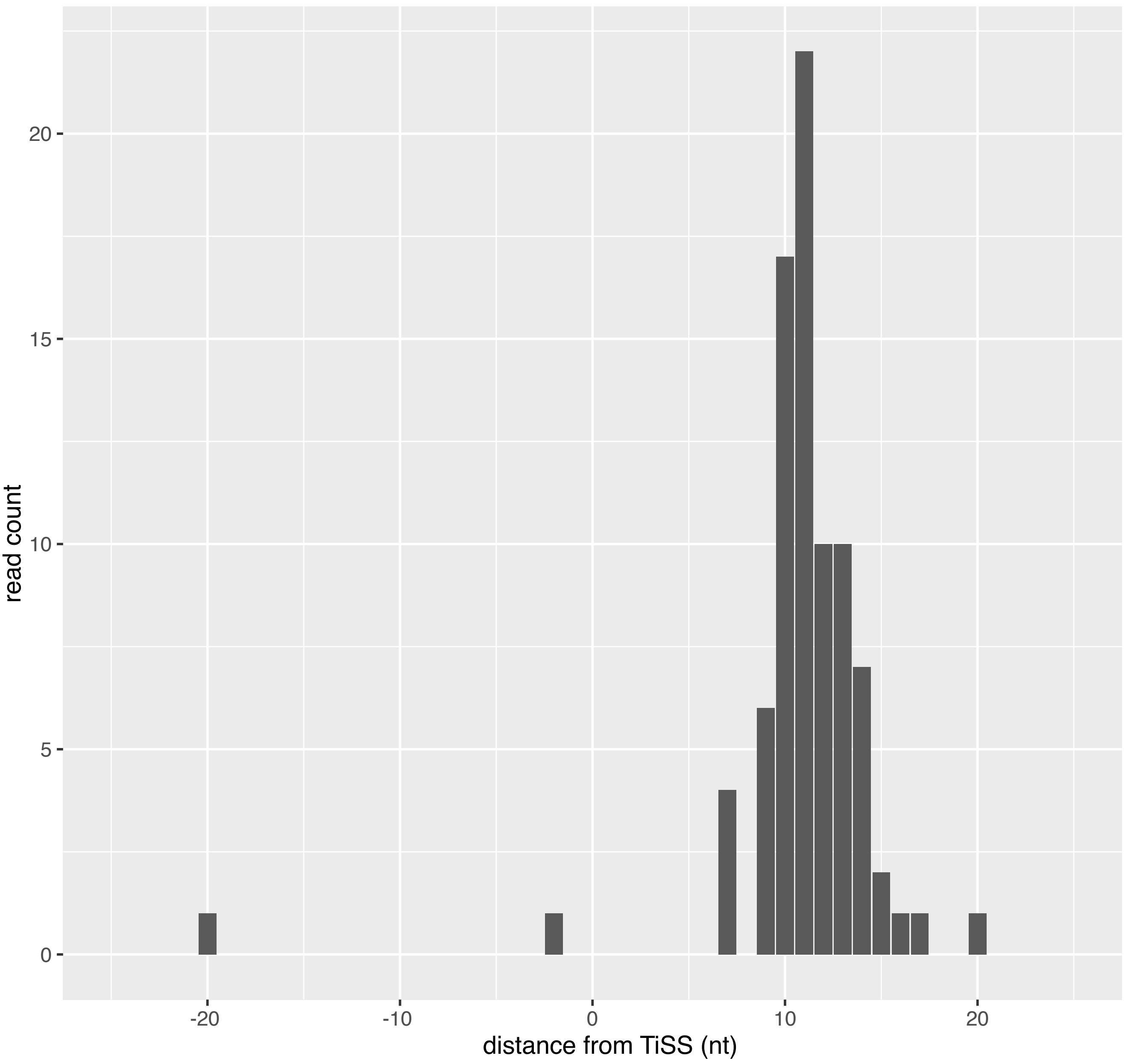

20
15
10
5
0
read count
-20
-10
0
10
20
distance from TiSS (nt)

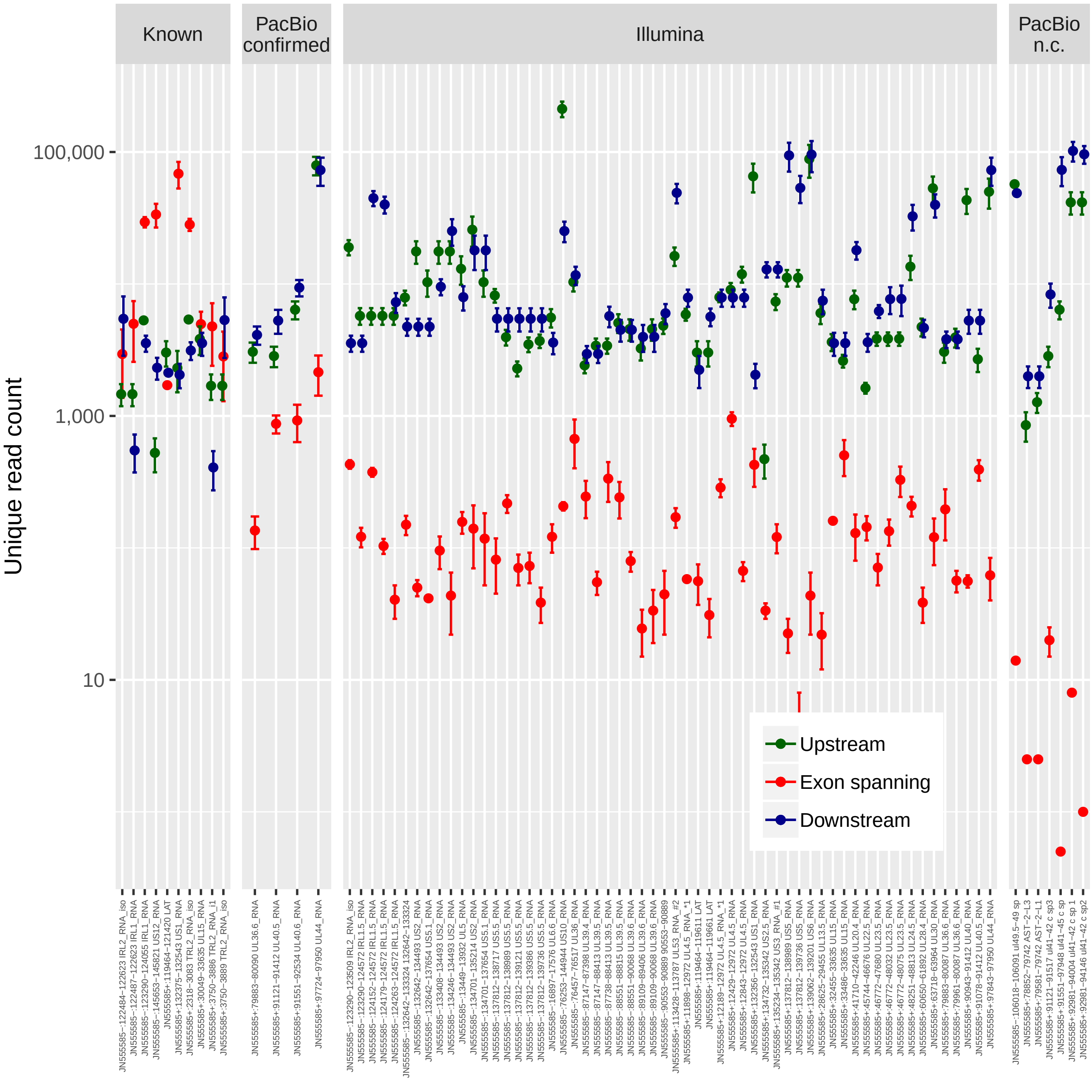
Known
PacBio confirmed
Illumina
PacBio n.c.
Unique read count
100,000
1,000
10
Upstream
Exon spanning
Downstream

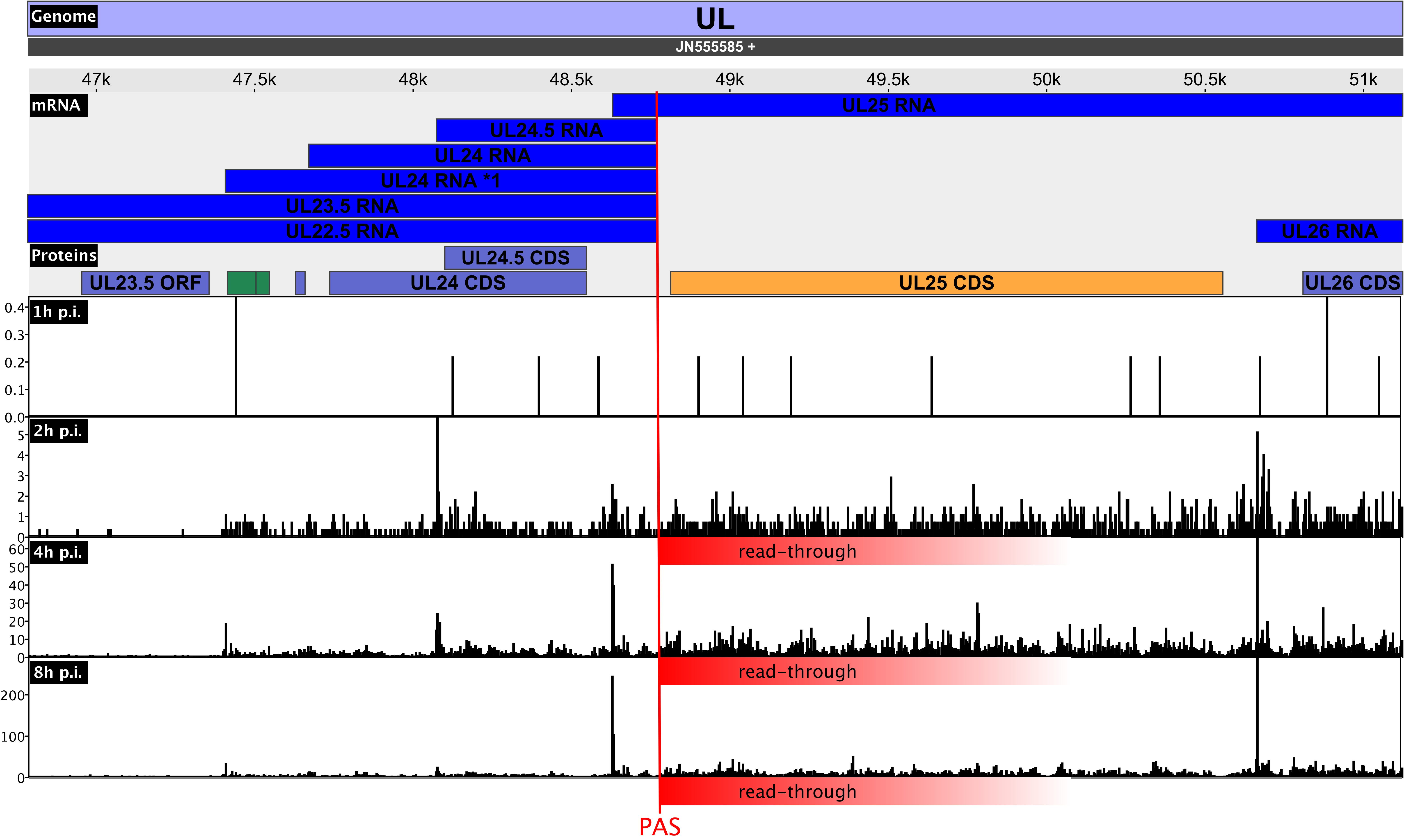

Genome
UL
JN555585 +
47k
47.5k
48k
48.5k
49k
49.5k
50k
50.5k
51k
mRNA
UL25 RNA
UL24.5 RNA
UL24 RNA
UL24 RNA *1
UL23.5 RNA
UL22.5 RNA
UL26 RNA
Proteins
UL24.5 CDS
UL23.5 ORF
UL24 CDS
UL25 CDS
UL26 CDS
1h p.i.
2h p.i.
4h p.i.
8h p.i.
read-through
read-through
read-through
PAS

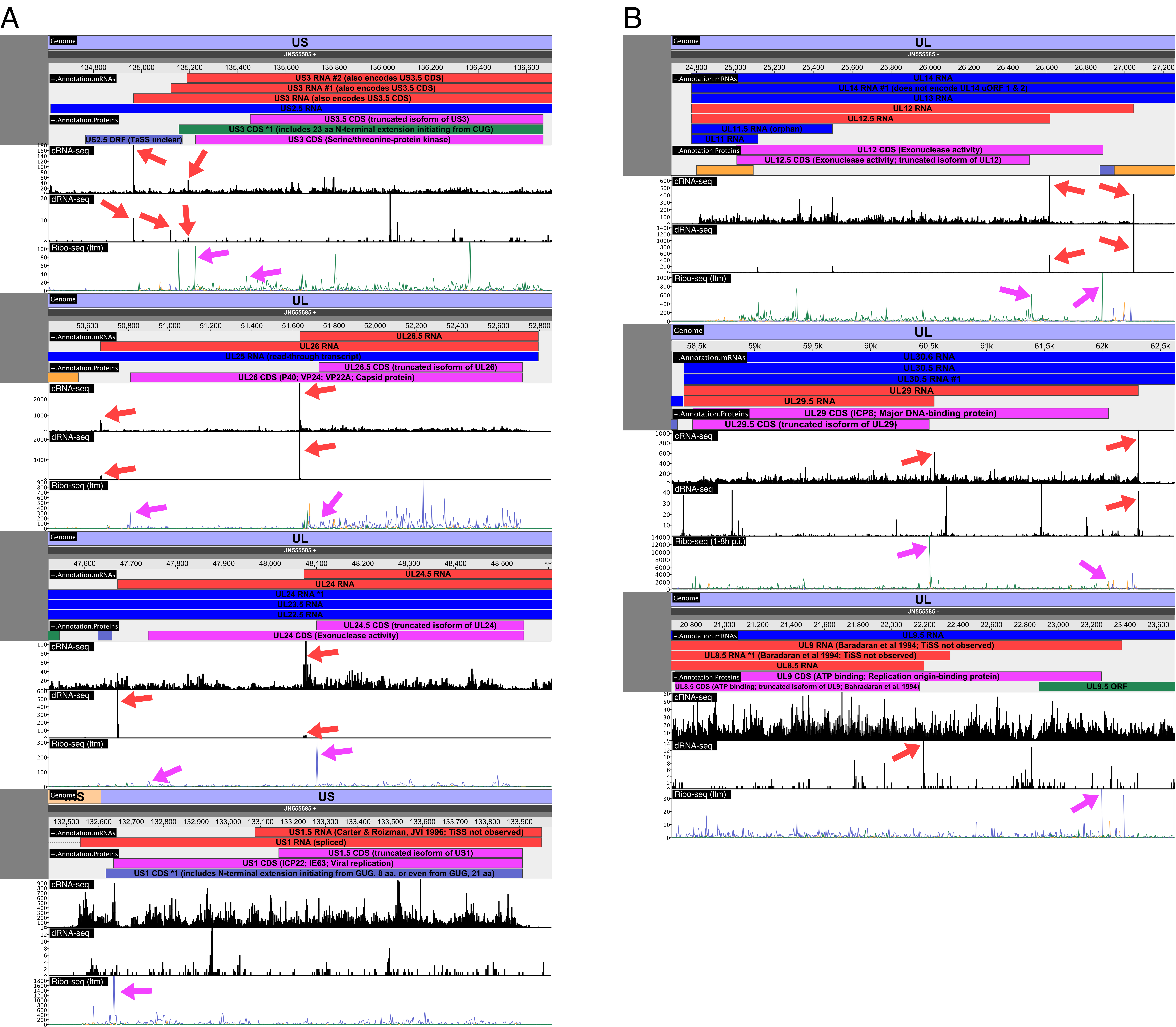
A
Genome
US
JN555585 +
+.Annotation.mRNAs
US3 RNA #2 (also encodes US3.5 CDS)
US3 RNA #1 (also encodes US3.5 CDS)
US3 RNA (also encodes US3.5 CDS)
US2.5 RNA
+.Annotation.Proteins
US3.5 CDS (truncated isoform of US3)
US3 CDS *1 (includes 23 aa N-terminal extension initiating from CUG)
US2.5 ORF (TaSS unclear)
US3 CDS (Serine/threonine-protein kinase)
cRNA-seq
dRNA-seq
Ribo-seq (ltm)
UL
UL26.5 RNA
UL26 RNA
UL25 RNA (read-through transcript)
UL26.5 CDS (truncated isoform of UL26)
UL26 CDS (P40; VP24; VP22A; Capsid protein)
UL24.5 RNA
UL24 RNA
UL24 RNA *1
UL23.5 RNA
UL22.5 RNA
UL24.5 CDS (truncated isoform of UL24)
UL24 CDS (Exonuclease activity)
US1.5 RNA (Carter & Roizman, JVI 1996; TiSS not observed)
US1 RNA (spliced)
US1.5 CDS (truncated isoform of US1)
US1 CDS (ICP22; IE63; Viral replication)
US1 CDS *1 (includes N-terminal extension initiating from GUG, 8 aa, or even from GUG, 21 aa)
B
JN555585 -
-.Annotation.mRNAs
UL14 RNA
UL14 RNA #1 (does not encode UL14 uORF 1 & 2)
UL13 RNA
UL12 RNA
UL12.5 RNA
UL11.5 RNA (orphan)
UL11 RNA
-.Annotation.Proteins
UL12 CDS (Exonuclease activity)
UL12.5 CDS (Exonuclease activity; truncated isoform of UL12)
UL30.6 RNA
UL30.5 RNA
UL30.5 RNA #1
UL29 RNA
UL29.5 RNA
UL29 CDS (ICP8; Major DNA-binding protein)
UL29.5 CDS (truncated isoform of UL29)
Ribo-seq (1-8h p.i.)
UL9.5 RNA
UL9 RNA (Baradaran et al 1994; TiSS not observed)
UL8.5 RNA *1 (Baradaran et al 1994; TiSS not observed)
UL8.5 RNA
UL9 CDS (ATP binding; Replication origin-binding protein)
UL8.5 CDS (ATP binding; truncated isoform of UL9; Bahradaran et al, 1994)
UL9.5 ORF

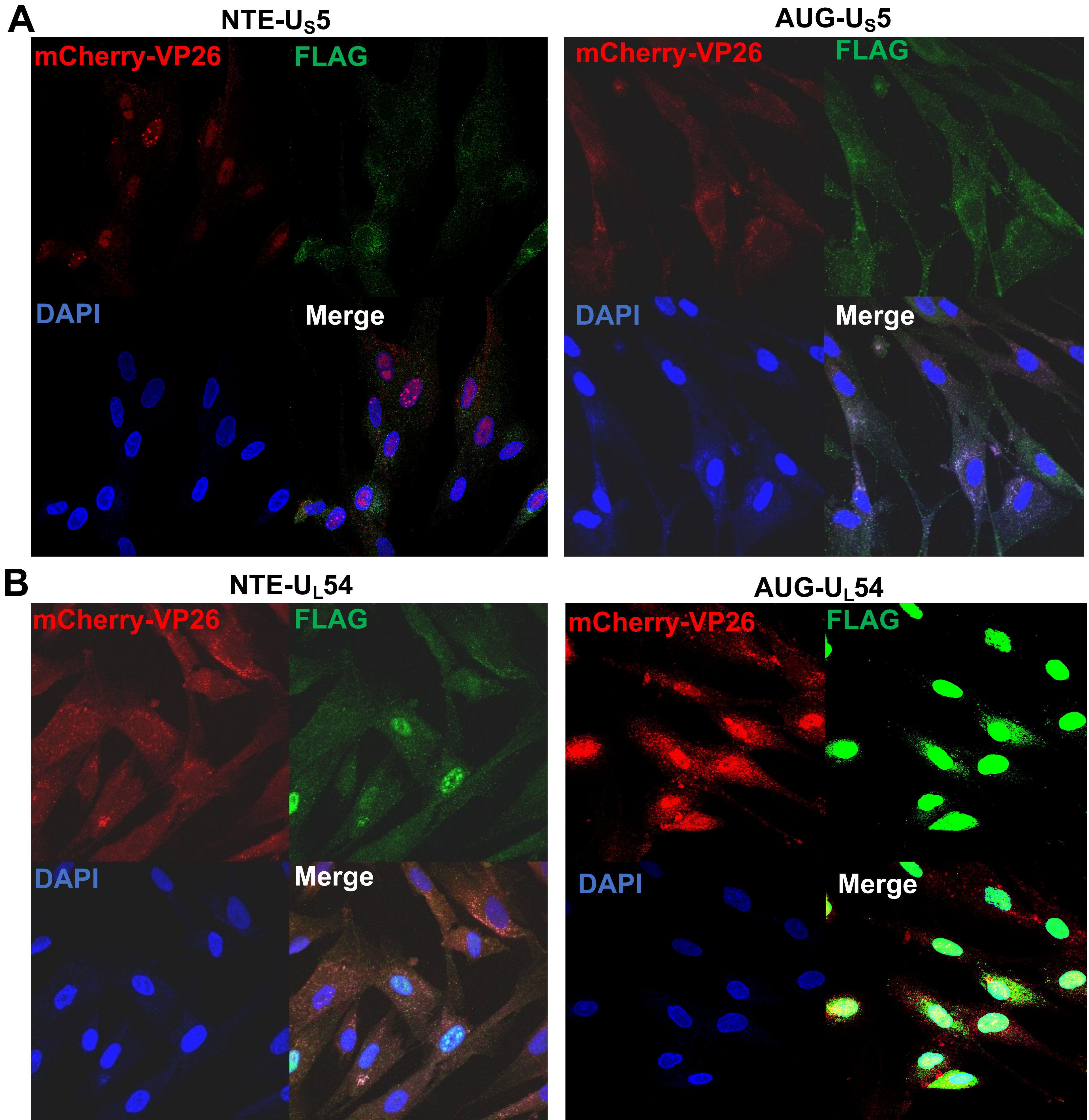
A
NTE-$U_S5$
AUG-$U_S5$
mCherry-VP26
FLAG
DAPI
Merge
B
NTE-$U_L54$
AUG-$U_L54$

| Virus | Accession number | Amino acid sequence |
| --- | --- | --- |
| HSV-1 | JN555585 | L/MPLLKTPGPVVRGARWLALTVRR**M** |
| HSV-2 | NC_001798 | L/MPLLKTPGPVARGARWLARATRQ**M** |
| BHV-1 | NC_001847 | IAGVDRVRLGVRLPFLPQARSRDTTRRSWAPM |
| FeHV-1 | NC_013590 | RFLLFRKCIRLANMDRFPRVGLSCCIPTSKGDIDTGDNYKLQST**M** |
| MaHV-1 | KT594769 | TCLSFQITGSLC**M** |
| PRV | NC_006151 | MLAMWRWVTKRSRLRRGHAHLGGNKGVRGICSLYLAGLSRGLSRVHAQRSHAAT**M** |

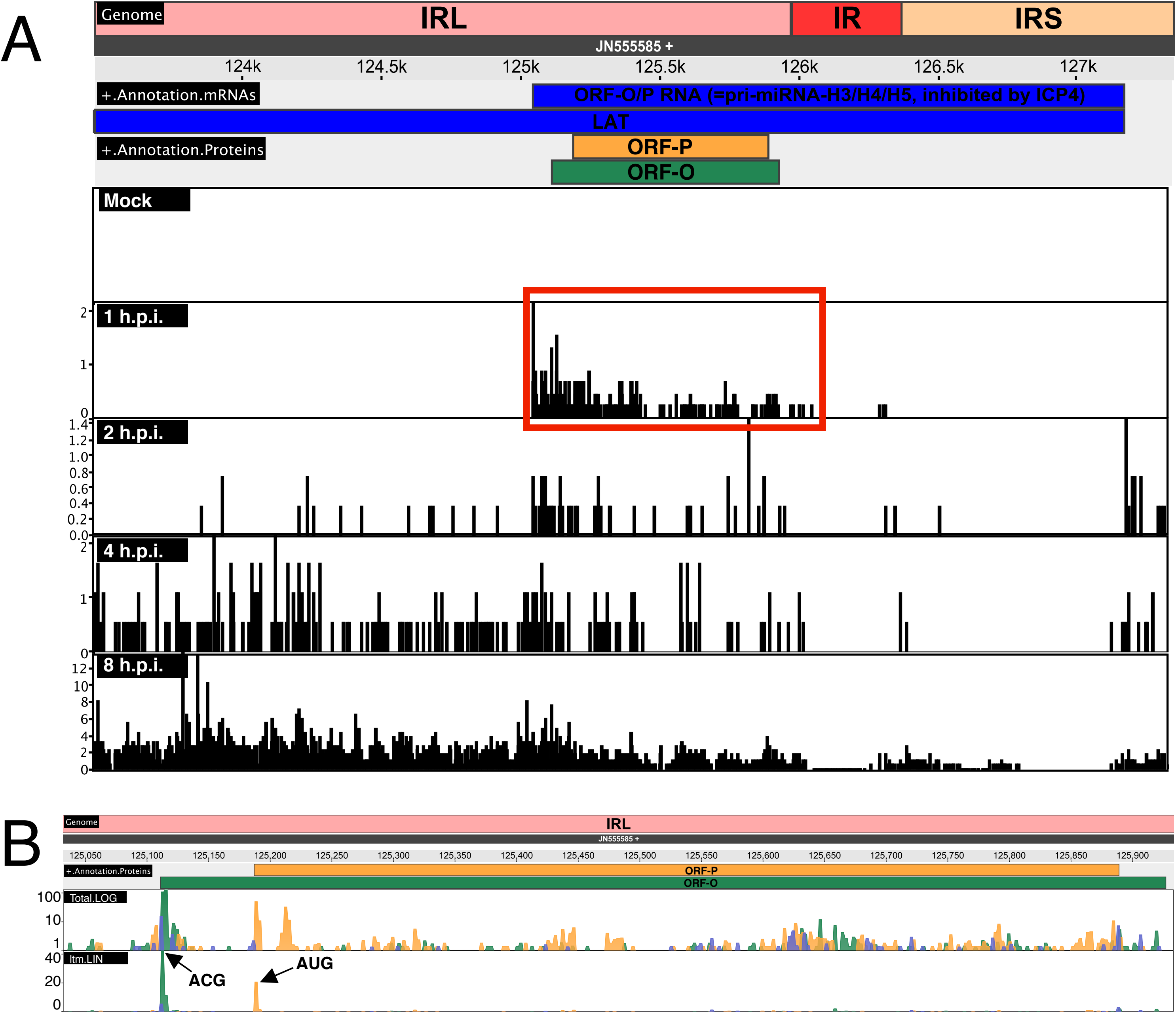
A
Genome
IRL
IR
IRS
JN555585 +
124k
124.5k
125k
125.5k
126k
126.5k
127k
+.Annotation.mRNAs
ORF-O/P RNA (=pri-miRNA-H3/H4/H5, inhibited by ICP4)
LAT
+.Annotation.Proteins
ORF-P
ORF-O
Mock
1 h.p.i.
2 h.p.i.
4 h.p.i.
8 h.p.i.
B
Genome
IRL
JN555585 +
125,050
125,100
125,150
125,200
125,250
125,300
125,350
125,400
125,450
125,500
125,550
125,600
125,650
125,700
125,750
125,800
125,850
125,900
+.Annotation.Proteins
ORF-P
ORF-O
Total.LOG
ltm.LIN
ACG
AUG

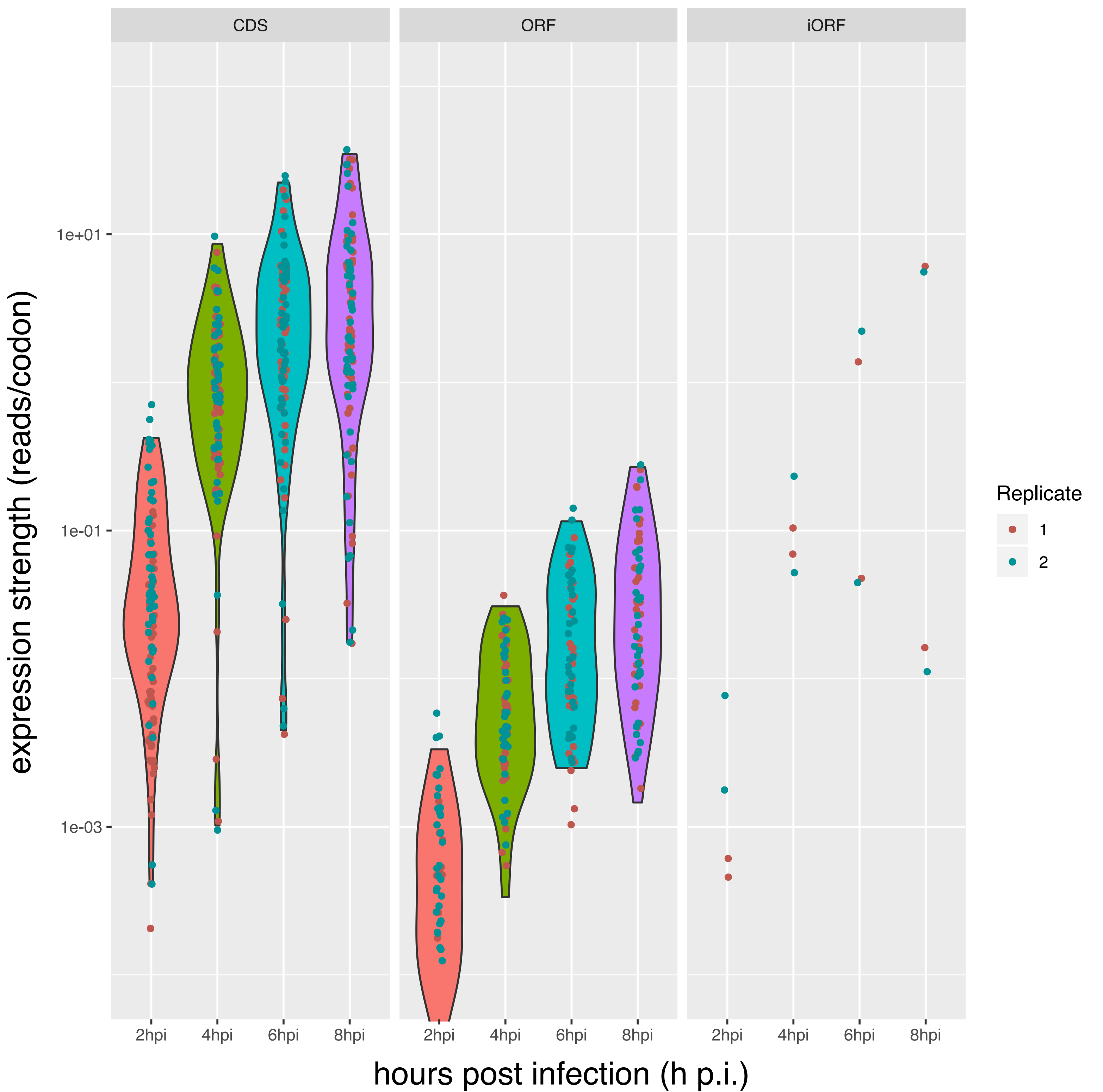

CDS
ORF
iORF
1e+01
1e-01
1e-03
expression strength (reads/codon)
2hpi
4hpi
6hpi
8hpi
hours post infection (h p.i.)
Replicate
1
2
