## Supplementary material for "Integrative functional genomics decodes herpes simplex virus 1": Principles of new nomenclature

**Principles of the new nomenclature of HSV-1 transcripts and ORFs**

1. No previously annotated ORFs were renamed to avoid causing confusions with previous work. All viral ORFs and transcript mentioned in the 6^th^ Edition of Fields of Virology were included.
2. To differentiate all new ORFs from the previously reported ORFs, we labeled all previous ORFs as “coding sequences”, e.g. UL1 CDS.
3. We differentiate long (≥100aa; named “ORF”) from short (3 – 99aa) viral ORFs.
4. We differentiated five different kinds of sORFs. These include upstream open reading frames (“uORFs”), upstream overlapping ORFs (“uoORFs”), internal ORFs (“iORFs”) and downstream ORFs (“dORFs”). In addition, sORFs, which are expressed from transcripts not containing any large ORF were named “sORFs”, e.g. UL34.5 sORF 1 and 2.
5. Translation of “uORFs” both starts and terminates upstream of a large ORF. A transcript can have multiple uORFs (e.g. UL14 uORF 1 and 2). In case a transcript does not encode any ORF >100aa, all short ORFs it encodes are labeled “sORFs”, e.g. UL30.5 sORF 1 and UL30.5 sORF 2.
6. In contrast to uORFs, uoORFs overlap with the main ORF expressed from the respective transcript.
7. Internal ORFs (iORFs) are located within the coding sequence of large ORFs but expressed in a different frame. In principle, two scenarios can explain their translation.
(i) They can be translated by ribosomes, which have missed the TaSS of the main ORF (e.g. UL20 iORF) and thus initiate translation at the iORF.
(ii) They can result from alternative independent transcripts initiating downstream of the respective TaSS of the main ORF, e.g. UL53 iORF RNA #2.

iORFs were thus not labeled as “orphan”.

8. Finally, a small number of downstream ORFs (dORFs) were annotated. These represent sORFs located downstream of large ORFs, which could not be explained by an independent transcript, e.g. UL39.6 dORF 1 and 2 downstream of UL39.6 ORF. Their translation may result from ribosomes re-initiating after completing the translation of the large ORF located further upstream. Therefore, they were not labeled as “orphan”. However, in most cases it is equally likely that they are translated from a yet unidentified viral transcript.
9. In principle, novel viral transcripts, ORFs and sORFs can all result in the introduction of a new viral gene identifier, e.g. UL28.5.
   a. Any novel large viral ORF, e.g. UL36.5 ORF, was given a new identifier unless it was overlapping with another large ORF. In the rare case that two overlapping viral ORFs (translated from different frames) were obviously expressed from the same transcript, these were named A and B, e.g. UL40.7A ORF and UL40.7B ORF as well as TRL2 CDS and TRL2A ORF.
   b. For viral transcripts to be given a new identifier, this required a transcription start site (TiSS) >500 nucleotides upstream of the closest other transcript, e.g. UL54.5 RNA (orphan).
   c. Any sORF >20aa in length that could not be attributed to another viral gene as a either uORF, uoORF, iORF or dORF was given a new identifier, e.g. UL27.5 sORF 1.
10. Numbering of new identifiers was defined based on the location of the TiSS or TaSS in relation to the neighboring previously annotated genes (x and x+1) on either strand. In case multiple new identifiers were required between two annotated genes, the most strongly expressed gene was named x.5, the neighboring ones x.4 and x.6. As annotations of additional genes by previous studies did not all follow the same rules in regards to neighboring genes, we tried to choose the best possible numbering for each locus.
11. Alternative transcription start sites is a very common phenomenon in the viral genome. Many of the additional transcriptions contain additional uORFs and thereby explain their expression. As such, we commonly observed >1 distinct TiSS within a window of 250 nt up- or downstream of the transcript of a given locus. The main TiSS was defined by the highest cRNA-seq or dRNA-seq peak. Within a window of +/-10 nt, no additional TiSS were annotated. TiSS identified by cRNA-seq, dRNA-seq and PacBio commonly matched perfectly at single nucleotide level.
12. Any transcript that did not contain an ORF within its first 500 nucleotides (nt) was labeled as “orphan”, e.g. UL54.5 RNA (orphan).
13. Any ORF or sORF for which no transcript could be identified that explained its translation within the transcript’s first 500 nt was labeled as “orphan”, e.g. US11.5 ORF (orphan).
14. Additional transcripts initiating upstream of the main transcript were labeled “*”+”number” with higher numbers reflecting increasing distance to the main TiSS, e.g. UL24 RNA *1.
15. Transcripts initiating downstream of the main transcript were labeled “#”+”number” with higher numbers reflecting increasing distance to the TiSS of the main transcript, e.g. UL41 RNA #1 and UL41 RNA #2.
16. Transcript experiencing alternative splicing were labeled as “iso1, iso2…”.
17. The annotation of uORFs was based on the most prominent transcript of the respective locus, e.g. UL6 uORF. Alternative TiSS commonly explained the expression of additional uORFs, e.g. UL6 RNA *1 explained UL6 uORF RNA *1.
18. Transcripts with retained introns were labeled as “i”+”number”, e.g. IRL2 RNA i1. The respective ORF variants were labeled accordingly, e.g. IRL2 ORF RNA i1
19. N-terminal extensions of ORFs were labeled with “*1”, e.g. UL50 CDS *1. In case of a second, longer N-terminal extension this was labeled “*2”, e.g. UL50 CDS *2. All N-terminal extensions of previously identified proteins initiated from non-AUG start codons. Both the start codon and the length of the extension are indicated in brackets, e.g. US3 CDS *1 (includes 23 aa N-terminal extension initiating from CUG).
20. N-terminal truncations of ORFs were labeled with “#”+”number”, e.g. UL37.6 ORF #1. We did not observe any more than 1 truncated version of a given ORF.
21. ORFs, sORFs and transcripts expressed from the repeat regions of the viral genome where named accordingly, e.g. IRL2.5 ORF and TRL2.5 ORF. We did not differentiate the three other possible orientations of the unique long and unique short regions.

**Annotations in the vicinity of the latency-associated transcript:**

We were unable to detect LAT in our data from lytically infected HFF but annotated it nevertheless. However, translation of the two previously described viral ORFs, ORF-O and ORF-P was readily detectable. In addition, we identified the corresponding RNA (ORF-O/P RNA). Both ORFs and the respective transcript were only apparent at early but not late times of infection. This is consistent with its reported repression by ICP4. However, our data indicate that ORF-O does not result from a frameshift in ORF-P but rather initiates from an ACG start codon 76nt upstream of the start codon of ORF-P.
